# eIF4E phosphorylation recruits β-catenin to mRNA cap and selectively promotes Wnt pathway translation in dentate gyrus LTP maintenance in vivo

**DOI:** 10.1101/2022.09.28.509312

**Authors:** Sudarshan Patil, Kleanthi Chalkiadaki, Tadiwos Feyissa Mergiya, Konstanze Simbriger, Inês S. Amorim, Shreeram Akerkar, Christos G. Gkogkas, Clive R. Bramham

**Author notes:** KS: Department of Pharmacology, Medical University Innsbruck, 6020, Innsbruck, Austria. Correspondence (C.R.B.), (S.P.), (C.G.G.).

## Abstract

The mRNA cap-binding protein, eukaryotic initiation factor 4E (eIF4E), is crucial for translation and regulated by Ser209 phosphorylation. However, the biochemical and physiological role of eIF4E phosphorylation in translational control of long-term synaptic plasticity is unknown. We demonstrate that phospho-ablated *Eif4e^S209A^* knockin mice are profoundly impaired in dentate gyrus LTP maintenance in vivo, while basal perforant path-evoked transmission and LTP induction are intact. mRNA cap-pulldown assays show that phosphorylation is required for synaptic activity-induced removal of translational repressors from eIF4E, allowing initiation complex formation. Using ribosome profiling, we identified selective, phospho-eIF4E-dependent translation of the Wnt signaling pathway in *in vivo* LTP. Surprisingly, the canonical Wnt effector, β-catenin, was massively recruited to the eIF4E cap complex following LTP induction in wild-type, but not *Eif4e^S209A^*, mice. These results demonstrate a critical role for activity-evoked eIF4E phosphorylation in dentate gyrus LTP maintenance, bidirectional remodeling of the mRNA cap-binding complex, and mRNA-specific translational control linked to Wnt pathway.

**Key highlights:**

1. Synaptic activity-induced eIF4E phosphorylation controls DG-LTP maintenance in vivo
2. eIF4E phosphorylation triggers release of translational repressors from cap complex
3. eIF4E phosphorylation recruits β-catenin to cap complex
4. eIF4E phosphorylation selectively enhances translation of Wnt pathway

## INTRODUCTION

Neuronal activity-dependent synaptic plasticity is crucial for adaptive behaviors such as memory formation (Josselyn and Tonegawa, 2020), and dysregulation of plasticity is commonly found in animal models of neurodevelopmental and degenerative disorders. At excitatory glutamatergic synapses, the generation of stable structural and functional changes lasting hours or more requires de novo gene expression and protein synthesis (Matthies et al., 1990; Nguyen and Kandel, 1996; Panja and Bramham, 2014). Global gene expression profiles in various plasticity models have been elucidated using microarrays and RNA-sequencing (Havik et al., 2007; Maag et al., 2015; Park et al., 2006; Ryan et al., 2011; Wibrand et al., 2006) as well as ribosome profiling (Chen et al., 2017; Hien et al., 2020). In neurons, regulation of gene expression at the level of translation is critical for long-term synaptic plasticity (Costa-Mattioli et al., 2009; Klann and Dever, 2004).

Translation initiation, the multistep process by which the ribosome is recruited to mRNA, is tightly regulated and often rate-limiting for protein synthesis (Sonenberg and Hinnebusch, 2009). Eukaryotic translation initiation factor 4E (eIF4E), which binds to the 5’-terminal cap structure of cytoplasmic mRNA, plays a key role in both the process and regulation of translation. eIF4E enables assembly of a multiprotein translation initiation complex at the mRNA 5 ’ end, which recruits the 40S small ribosomal subunit and scans to the mRNA start codon. Interaction of eIF4E with the scaffolding protein eIF4G is crucial for initiation complex formation. This interaction is obstructed by eIF4E-binding proteins (4EBPs). Hypophopshorylated 4E-BPs bind to eIF4E, competing for eIF4G binding and repress translation initiation. Signaling to the kinase mechanistic target of rapamycin (mTORC1) enhances translation by phosphorylation of 4E-BPs, leading to their dissociation from eIF4E (Gingras et al., 2001; Proud, 2009). In a major convergent pathway, activation of extracellular signal-regulated kinase (ERK, aka mitogen-activated protein kinase; MAPK) signaling to MAPK-interacting kinases (MNK1 and MNK2) phosphorylates eIF4E on a single residue, Ser209 (Wang et al., 1998; Waskiewicz et al., 1997). Phosphorylation of eIF4E is usually, but not always, associated with enhanced translation initiation (Beggs et al., 2015; Bramham et al., 2016; Joshi, 2014; Kong and Lasko, 2012). Thus, the molecular function of Ser209 eIF4E phosphorylation is unresolved (Chalkiadaki et al., 2021).

Translational control in synaptic plasticity has been extensively studied in excitatory pathways of the hippocampus. A major question is whether translation mechanisms are differentially implemented to sculpt protein synthesis and plasticity in a pathway-specific manner. In the hippocampal CA1 region, ERK-dependent translation initiation regulates stable LTP formation (Gelinas et al., 2007; Kelleher et al., 2004). Surprisingly, eIF4E phosphorylation is dispensable at Schaffer collateral-CA1 synapses, as LTP induction and maintenance are normal in knockin mice harboring nonphosphorylatable Ser209Ala mutated eIF4E (*Eif4e^S209A^* or *Eif4e^ki^*) (Amorim et al., 2018). At the medial perforant path input to the dentate gyrus (DG), studies employing kinase inhibitors and MNK knockout mice implicate ERK-MNK signaling in enhanced translation initiation and LTP maintenance (Panja et al., 2009, Panja et al., 2014). However, the specific role of eIF4E phosphorylation in DG-LTP maintenance and translation control has not been investigated.

Here, we used *Eif4e^ki^* mice to explore the role of eIF4E phosphorylation in DG-LTP in vivo. We demonstrate that ablation of Ser209 phosphorylation inhibits the synaptic activity-evoked discharge of eIF4E repressor proteins, prevents formation of the translation initiation complex, and selectively inhibits LTP maintenance without affecting basal synaptic transmission or LTP induction. Using unbiased ribosome profiling, we show that eIF4E phosphorylation is required for specific translation of Wnt family targets during DG-LTP. In canonical Wnt signaling, β-catenin mediates transcriptional regulation (Budnik and Salinas, 2011; Nusse and Clevers, 2017). Here we report a previously unknown, massive recruitment of β-catenin to eIF4E following LTP induction. The discharge of repressors and recruitment of β-catenin both depend on synaptic activity-evoked phosphorylation of eIF4E. Collectively, these results demonstrate a major function for Ser209 phosphorylation in stimulus-induced remodeling of eIF4E interactions and enhanced Wnt pathway translation in LTP maintenance.

## RESULTS

### Loss of eIF4E Ser209 phosphorylation selectively inhibits DG-LTP maintenance

We first examined the impact of ablating phospho-eIF4E on basal perforant path-DG transmission in adult anesthetized mice. Evoked field potentials were obtained across a range of stimulation intensities and input-output curves were constructed of the field EPSP slope (**Figure S1**A), population spike amplitude (**Figure S1**B), and plots of the EPSP-population spike relationship were made to evaluate synaptic excitability of granule cells (**Figure S1**C). Homozygous *Eif4e*^ki/ki^ mice were not significantly different from wild-type in any of these measures, indicating that ablation of phospho-eIF4E does not affect synaptic efficacy or granule cell excitability.

We then asked whether ablation of eIF4E phosphorylation impacts LTP induced by application of high-frequency stimulation (200 Hz, 4 trains of 15 pulses) (**Figure 1**). In wildtype mice, HFS induced an increase in fEPSP slope which remained stable during 3 hours of post-HFS recording (**Figure 1**A). In *Eif4e*^ki/ki^ mice, HFS induced an initial increase in fEPSP slope that was not significantly different in magnitude from wild-type control at 0-10 min post-HFS (**Figure 1**A and 1B). However, in striking contrast to wild-type, the fEPSP increase declined completely to baseline by 2 hours post-HFS. Already at 30-40 min post-HFS, *Eif4e*^ki/ki^ mice exhibited a severe (60.6 %) reduction in fEPSP potentiation relative to wildtype (**Figure 1**B and 1C). Like homozygotes, heterozygous *Eif4e*^ki/+^ mice showed impaired LTP at 30-40 min post-HFS (**Figure 1**B and 1C). However, the inhibition in heterozygotes was transient, as fEPSP responses returned to the wild-type control level to exhibit stable LTP (**Figure 1**A and 1B). The results show that phosphorylation of eIF4E is required for maintenance of synaptic LTP, but dispensable for basal transmission and LTP induction. Interestingly, *Eif4e*^ki/ki^ mice showed a stable increase in the population spike, indicating that mechanisms other than eIF4E phosphorylation regulate plasticity of granule cell excitability (**Figure 1**C, **Figure S2**A and 2B).

**Figure 1.**
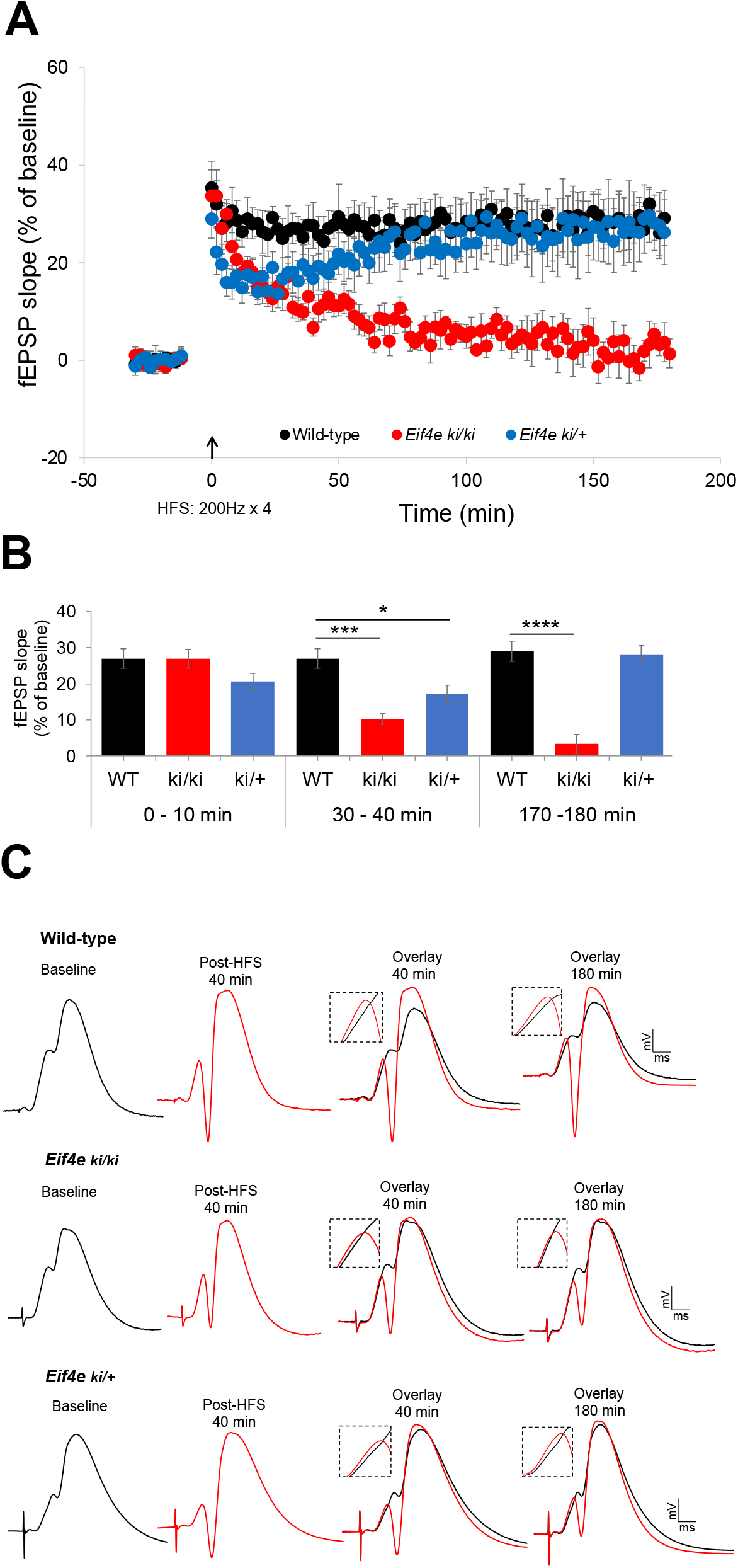
Ablation of phospho-eIF4E selectively impairs DG-LTP maintenance in vivo. (A) Time-course plots of medial perforant path-dentate gyrus (DG) evoked fEPSPs recorded before and after high-frequency stimulation (HFS, indicated by arrow) in homozygous *Eif4e*^S209A^ knockin mice (*Eif4e*^ki/ki^; n=7), heterozygous mice (*Eif4e*^+/ki^; n=8) and wild-type littermates (*Eif4e^+/+^* mice; n=10). Values are mean ± SEM of the maximum fEPSP slope expressed in percent of baseline. (B) Bar graphs show mean changes in fEPSP slope recorded between 0-10 min, 30-40 min, and 170-180 min post-HFS. Significant differences between genotypes: *p < 0.05; ***p < 0.0001, ****p < 0.00001; Student’s t-test. Heterozygotes exhibit transient impairment in early LTP maintenance. Homozygous show loss of stable LTP maintenance. (C) Representative field potentials recorded at baseline and post-HFS (40 min and 180 min). Each trace is the average of 4 consecutive responses. See also Figures S1 and S2.

### Loss of eIF4E phosphorylation inhibits stimulus-induced release of eIF4E repressors and prevents initiation complex formation in DG-LTP in vivo

Next, we examined the biochemical function of eIF4E phosphorylation. Previous work showed that pharmacological inhibition of MNK prevents eIF4E phosphorylation and formation of the translation initiation complex during DG-LTP. According to this model, MNK activity triggers discharge of eIF4E repressors: cytoplasmic FMRP interacting protein-1 (CYFIP1) and the canonical eIF4E-binding protein (4E-BP2) (Panja et al., 2009, 2014) (**Figure 2**A). However, the specific role of eIF4E phosphorylation is unknown, as MNK has multiple additional substrates with roles in translation and mRNA metabolism (Bramham et al., 2016; Buxadé et al., 2005; Joshi, 2014).

**Figure 2.**
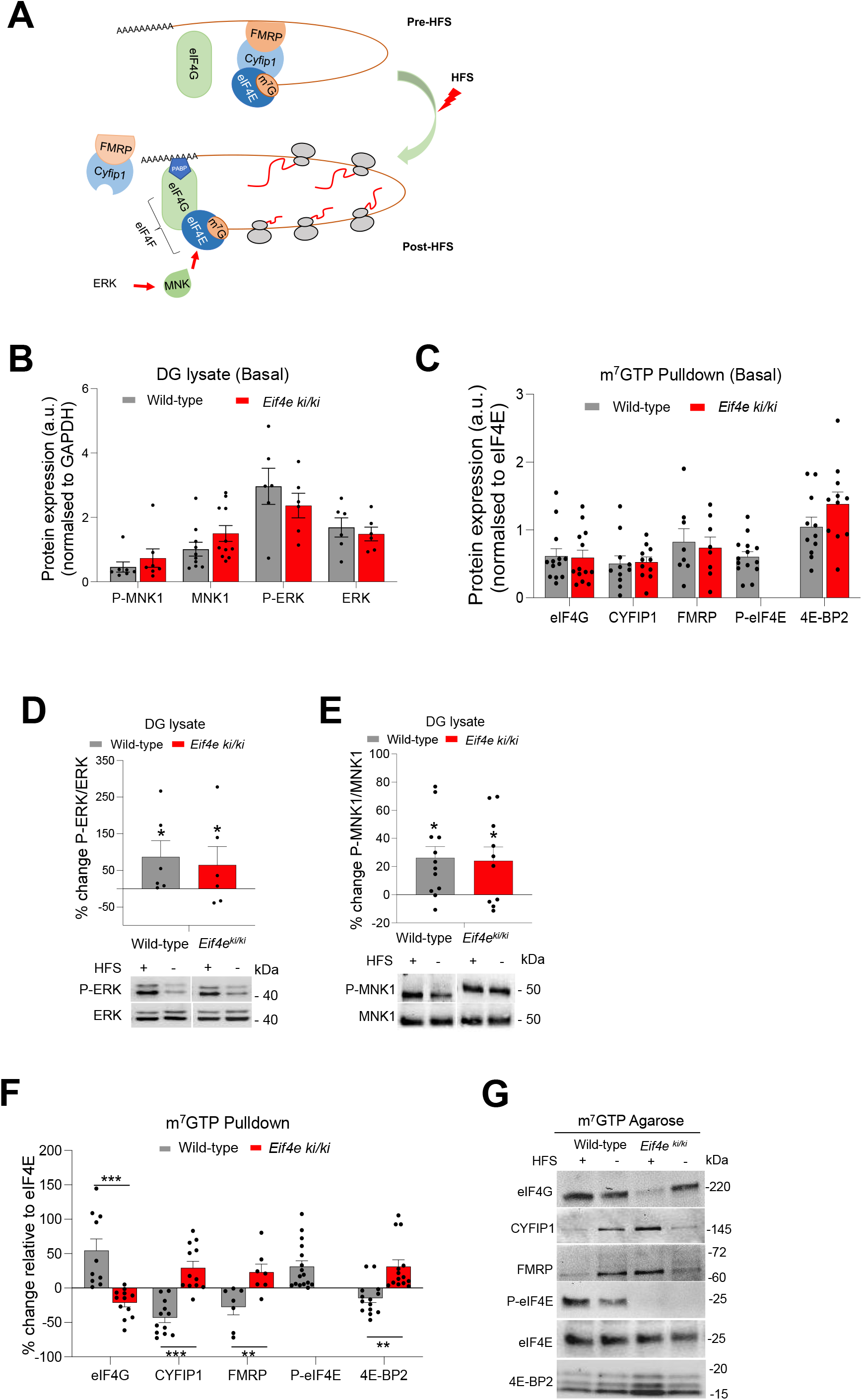
Ablation of phospho-eIF4E impairs synaptic activity-evoked discharge of eIF4E repressors and assembly of translation initiation complex. (A) Model of HFS-evoked, MNK1-catalyzed eIF4E phosphorylation resulting in discharge of the CYFIP1/FMRP (Fragile-x messenger ribonucleoprotein) complex from eIF4E and recruitment of eIF4G to form the translation initiation complex. Based on Panja et al. (2014). (B) Immunoblot analysis of total (T) and phosphorylated (P) ERK and MNK in DG lysates from naïve mice (basal state). No significant difference between wild-type and *Eif4e*^ki/ki^ mice in expression of active, P-ERK (Thr202/Tyr204), P-MNK (Thr197/202), or expression of their respective total proteins normalized to GAPDH. (P-ERK, n=6; ERK, n=6; P-MNK1, n=7; MNK1, n=11) Values are means + SEM. (C) Immunoblot analysis from m^7^GTP pulldown assays in DG lysates in naïve mice (basal state). Densitometric values for eIF4E cap-associated proteins (eIF4G, 4E-BP2, CYFIP1 and FMRP) are normalized to values of m^7^GTP-bound eIF4E. Ablation of phospho-eIF4E in *Eif4e*^ki/ki^ mice did not alter translation factor binding relative to wild-type. (D) (Top). Enhanced ERK phosphorylation in ipsilateral, HFS-treated DG. Values in lysates sample are expressed in percent change relative to contralateral DG control. P-ERK is normalized to total ERK. Upper and lower bands (ERK1/ERK2) combined for quantification. (Bottom). Representative immunoblots. *p < 0.05; Student’s t-test. (E) (Top). Enhanced MNK1 phosphorylation in ipsilateral, HFS-treated DG. Values in lysates sample are expressed in percent change relative to contralateral DG control. P-MNK1 is normalized to total MNK1. (Bottom). Representative immunoblots. HFS = high-frequency stimulation. (+) Ipsilateral DG, (-) Contralateral DG. In panels D and E, * indicates significant increase in HFS-treated DG relative to contralateral control. *p < 0.05; Student’s t-test. (F) Immunoblot analysis of m^7^GTP pulldowns from DG lysates obtained 40 min post-HFS. Values normalized to levels of m^7^GTP-bound eIF4E and expressed in percent changes in HFS-treated DG relative to contralateral control. eIF4G = 0.00017, CYFIP1= 0.00004, FMRP= 0.008, T-BP2= 0.0005 (**p < 0.001, ***p < 0.0001; Multiple t-test). (G) Representative immunoblots for panel F. See also Figure S3.

First, we asked whether ablation of Ser209 eIF4E phosphorylation impacts basal ERK-MNK signaling and initiation complex formation. In unstimulated DG tissue from naïve mice, expression of total and phosphorylated (activated) ERK and MNK did not significantly differ between *Eif4e*^ki/ki^ mice and wild-type (**Figure S2**B). eIF4E binds to the 7-methylguanosine (m^7^GTP) moiety of the 5’-terminal mRNA cap. To assess regulation of the eIF4E cap-binding complex, m^7^GTP cap-pulldowns were performed in DG lysates and immunoblotting was used to quantify levels of interacting proteins relative to eIF4E. Phospho-eIF4E was present in cap-pulldown samples from wild-type mice, but absent from *Eif4e*^ki/ki^ mice. However, there was no difference between genotypes in eIF4G binding to eIF4E, indicating normal formation of the eIF4F initiation complex (**Figure 2**C). Levels of cap/eIF4E-associated CYFIP1, FMRP, and 4E-BP2 in *Eif4e*^ki/ki^ mice were also not different from wild-type (**Figure 2**C). Immunoblotting of DG lysate (input) samples similarly showed no genotype differences in expression of the translation factors relative to GAPDH loading control (**Figure S3**A). Thus, under basal conditions, absence of eIF4E phosphorylation did not alter ERK-MNK signaling or eIF4E protein-protein interactions.

Next, we analyzed ERK-MNK signaling in response to perforant path stimulation. At 40 min post-HFS, expression of phospho-ERK and phospho-MNK was significantly enhanced in ipsilateral DG relative to the contralateral, non-stimulated DG, with no significant difference between genotypes (**Figure 2**D and 2E). Thus, HFS-induced ERK-MNK signaling is intact and unaltered in *Eif4e*^ki/ki^ mice. We then performed cap-pulldown assays in DG lysates to assess changes in eIF4E interactions. In wild-type mice, binding of CYFIP1, FMRP, and 4E-BP2 to eIF4E was significantly reduced while loading of eIF4G was enhanced, relative to the contralateral control DG (**Figure 2**F and 2G). In contrast, HFS in *Eif4e*^ki/ki^ mice failed to discharge the repressor proteins or enhance eIF4G binding to eIF4E (**Figure 2**F and 2G). Rather, we observed increased recovery of CYFIP1 and 4E-BP2 in the cap-pulldown in *Eif4e*^ki/ki^ mice, suggesting stabilization of the interaction complex in the absence of eIF4E phosphorylation. In DG lysates, CYFIP1 levels decreased in HFS-treated wild-type mice and increased in *Eif4e*^ki/ki^ mice, consistent with degradation of CYFIP1 after release from eIF4E (**Figure S3**B and 3C). Taken together, these results suggest that stimulus-evoked phosphorylation of eIF4E on Ser209 is required for the release of translational repressors, formation of the translation initiation complex and maintenance of LTP.

Translation of the immediate early gene Arc is causally implicated in DG-LTP consolidation (Messaoudi et al., 2007), and regulated by MNK signaling in rats and mice (Gal-Ben-Ari et al., 2012; Panja et al., 2009). In the present study, HFS-induced Arc expression was significantly reduced in *Eif4e*^ki/ki^ mice relative to wild-type (**Figure S3**B and 3C), whereas basal Arc expression did not differ between genotypes (**Figure S3**A). Taken together these data suggest that Arc expression in LTP specifically depends on MNK catalyzed phosphorylation of eIF4E.

### Ribosome profiling identifies phospho-eIF4E-dependent specific translation in LTP: enhanced translation of Wnt signaling pathway

Next, we used unbiased ribosome profiling to identify changes in translational activity linked to phospho-eIF4E-dependent maintenance of LTP. This analysis focused on 40 min post-HFS as a critical time in the transition to stable LTP. In the LTP experiments, mice received HFS along with the standard low-frequency test-pulse stimulation (LFS) to assess changes in the fEPSP. The ipsilateral, HFS-treated DG and the contralateral unstimulated DG were collected at 40 min post-HFS) (**Figure 3**A). To control for effects of test-pulse stimulation, a control group received LFS only) (**Figure 3**A). To ascertain the effect of HFS, we normalized the data from ipsilateral HFS-treated DG to LFS-treated DG from respective WT and *Eif4e*^ki/ki^ mice. We prepared RNA sequencing libraries from both ribosome-protected footprints (a proxy for translation) and total mRNA (a proxy for transcription) (**Figure 3**A). Novaseq produced high quality reads for footprints and mRNA libraries because: i) the distribution of footprint size (28-32nt) is canonical (**Figure S4**B, top panel), ii) the read distribution within the three frames is more abundant for the protein coding frame (**Figure S4**B, bottom panel), and iii) the periodicity of ribosomal footprints across mRNA coding and non-coding regions is canonical (**Figure S4**C).

**Figure 3.**
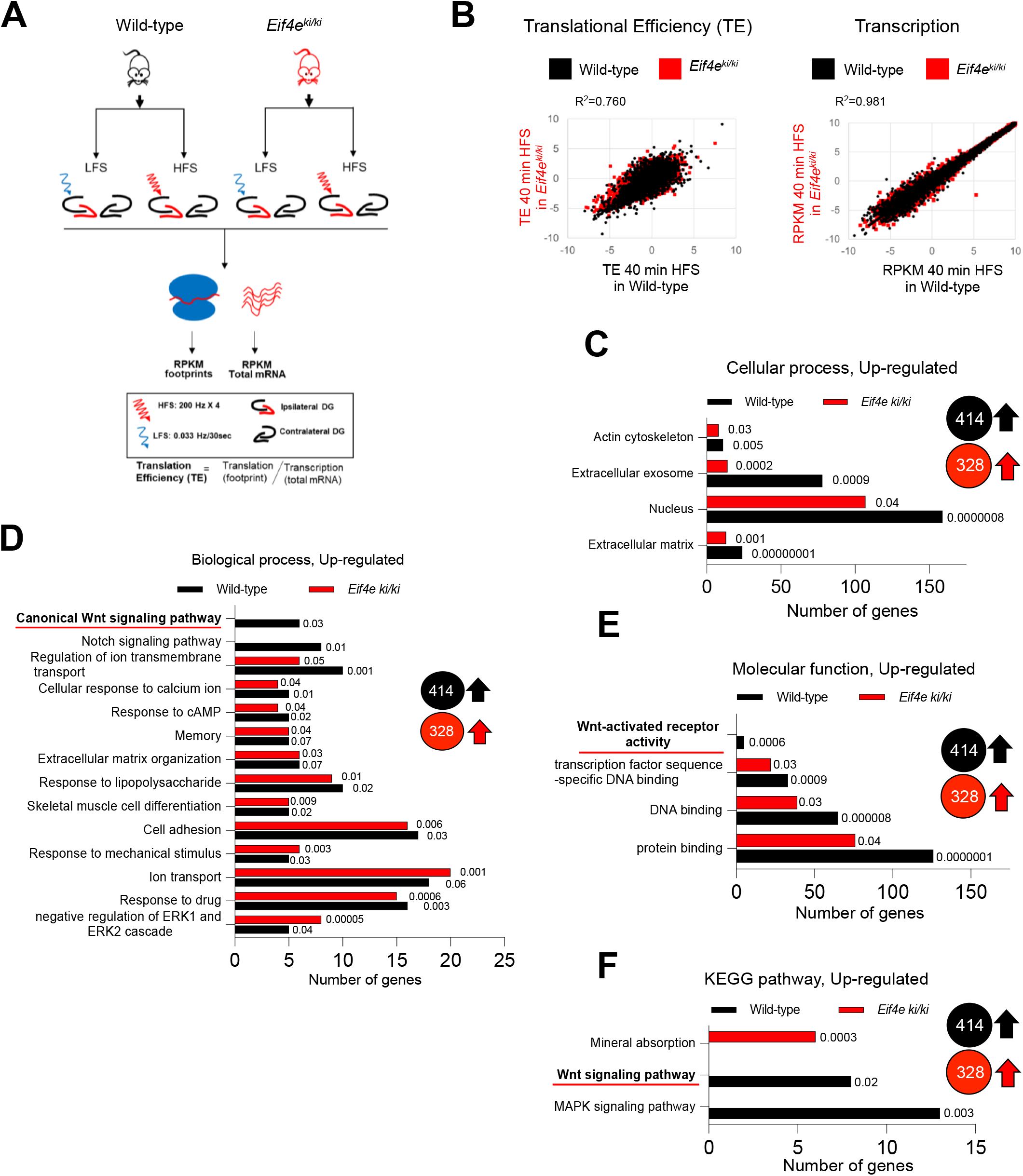
Ribosomal profiling identifies phospho-eIF4E-dependent specific translation of Wnt family pathway in DG-LTP in vivo. (A) Schematic representation of plan of ribosomal profiling experiment. (B) Scatter plot of log2 TE Plot showing upregulated translational efficiency and transcription in *Eif4e*^ki/ki^ mice versus *Eif4e^+/+^* (wild-type) libraries following HFS stimulation for 40 min (p<0.05 and 0.75 ≥TE ratio≤1.5). Gene ontology analysis of upregulated genes in 40 min HFS *Eif4e^+/+^* (414) and *Eif4e*^ki/ki^ mice (328); plots for cellular component (C), biological process (D) and molecular function (E) with number of genes in each category with p-values. (F) KEGG pathway analysis for upregulated genes. Mean ± SEM and Student’s *t*-test in **Table S2**. LFS: low-frequency test pulse stimulation only, HFS: high-frequency stimulation, DG: Dentate Gyrus. See also Figures S5 and S6.

First, we aimed to elucidate the translational landscape of LTP in wild-type (*Eif4e*^+/+^) mice 40-min after HFS. In terms of global mRNA translation (TE was calculated by the RPKM reads of footprints normalized to mRNA abundance) and global mRNA abundance, there was no significant effect of HFS treatment relative to contralateral DG or LFS-treated DG (**Figure S5**; **Table S3).** However, we detected an overall increase in mRNA-specific translation at 40 min post-HFS of differentially translated genes (DTGs) and an overall increase in transcription of differentially expressed genes (DEGs), including known immediate early genes (IEGs) such as Arc, Junb, Npas4, Fos and Fosb (**Figure S5** and **Table S3)**. Gene ontology analysis of DTGs and DEGs using DAVID, and Ingenuity pathway analysis (IPA) revealed LTP-related categories, such as calcium, post-synapse, excitatory post-synapse potential and key biological pathways, such as AMPK signaling, actin cytoskeleton and extracellular matrix (**Figure S5** and **Table S4**).

Second, we investigated the effect of the ablation of Ser209 phosphorylation, focusing on translational efficiency (TE) at 40 min post-HFS. TE was calculated by the RPKM reads of footprints normalized to mRNA abundance. While we did not detect significant changes in global mRNA levels between HFS-treated wild-type and *Eif4e*^ki/ki^ (**Figure 3B**, R^2^=0.981), there was a modest upregulation of translation (**Figure 3B**, R^2^=0.760). Consequently, analysis of log_2_ of TE between wild-type and *Eif4e*^ki/ki^ mice at 40 min post-HFS, as compared to LFS-treated control (ratio<0.667&ratio>1.5; p<0.05), identified that 471 genes in wild-type mice (414 upregulated and 57 downregulated) and 419 genes in *Eif4e*^ki/ki^ mice (328 upregulated and 91 downregulated) were differentially translated (**Table S5**). We then performed GO analysis using the DAVID (database for annotation, visualization, and integrated discovery) platform (Huang et al., 2009) (**Figure 3** and **Table S4**). Key GO categories identified for upregulated DTGs both in wild-type and *Eif4e*^ki/ki^ mice include memory, extracellular matrix organization, cell adhesion, cellular response to calcium ion and actin cytoskeleton (**Figure 3C** and **3D, Table S4**). Strikingly, the canonical Wnt signaling pathway and its related genes were absent from the *Eif4e*^ki/ki^ mice GO analysis, whereas they were significantly upregulated in wild-type mice (**Figure 3D-3F, Table S4**). Within this GO Wnt signaling pathway category, we identified translationally upregulated Wnt4, Wnt receptors (Lrp5, Fzd2, Fzd4), and the receptor scaffolding protein dishevelled 2 (Dvl2) (**Table S6**). In addition, major β-catenin transcriptional targets (Adamts10, Bcl2l2, Ppard, Vegf, Hspg2, pkd1) exhibited enhanced translational efficiency at 40 min post-HFS relative to controls mice given LFS only (**Table S6**). Key GO categories among downregulated DTGs in *Eif4e*^ki/ki^ mice, as compared to wild-type, include protein synthesis, transcription, poly(A) RNA binding and ribosome (**Figure S4**).

The list of mRNAs identified in ribosomal footprinting in *Eif4e^+/+^* and *Eif4e*^ki/ki^ mice is shown in **Table S5.** Many Wnt pathway components have long and structured 5’ UTRs (Untranslated Regions) (Nguyen et al., 2018). Given the pronounced change in Wnt pathway mRNA translation, we further analyzed DTGs at 40 min post-HFS in the *Eif4e*^ki/ki^ vs *Eif4e^+/+^* mice, focusing on 5’ UTR mediated mechanisms. We find that the mRNAs of DTGs upregulated in *Eif4e^+/+^* mice at 40 min post-HFS harbor 5’ UTRs which are significantly longer, and more complex (higher %GC and high folding free energy) and are enriched in uORF, TOP and PG4 motifs compared with downregulated DTG (**Figure S7**). Remarkably, HFS in *Eif4e*^ki/ki^ mice did not elicit the same response as *Eif4e^+/+^* mice, displaying a loss of function phenotype with no significant differences neither in length, %GC, folding free energy nor in the incidence of uORF, TOP and PG4 motifs in upregulated DTG compared with downregulated (**Figure S7**). Taken together, these data suggest that eIF4E Ser209 phosphorylation is required in the DG for mRNA-specific translation during *in vivo* LTP and a major GO category regulated downstream is the Wnt signaling pathway.

### Synaptic activity-evoked eIF4E phosphorylation recruits atypical β-catenin to the translation initiation complex

In canonical Wnt signaling, β-catenin accumulates in the nucleus and activates transcription of Wnt target genes (Budnik and Salinas, 2011; Nusse and Clevers, 2017). However, a study in vascular smooth muscle cell cultures shows interaction of β-catenin with FMRP in the eIF4E cap-binding complex (Ehyai et al., 2018). In these cells, Wnt signaling triggers release of β-catenin from the complex, resulting in derepression of translation along with nuclear accumulation of β-catenin.

We aimed to determine whether β-catenin is part of the eIF4E-cap complex of adult DG, and, if so, whether it is regulated by LTP-inducing stimuli and Ser209 eIF4E phosphorylation. In DG from naïve unstimulated mice, β-catenin was detected in m^7^GTP cap-pulldowns and lysate samples, with no significant difference in expression between *Eif4e*^ki/ki^ and wild-type mice (**Figure 4**A-4C). Thus, under basal conditions, β-catenin associates with eIF4E in a manner that does not depend on eIF4E phosphorylation. Following LTP induction, β-catenin in lysate samples from HFS-treated DG was increased 20% relative to the contralateral control, but there was no difference between genotypes (**Figure 4**D and 4F). In contrast in the cap-pulldown assays, HFS in wild-type mice elicited a significant mean 78.3% enhancement of β-catenin levels while no change was found in *Eif4e*^ki/ki^ mice (**Figure 4**E and 4F). These data demonstrate recruitment of β-catenin to eIF4E that is evoked by HFS of perforant path synapses, dependent on eIF4E phosphorylation, and functionally linked to LTP maintenance and enhanced translation of Wnt signaling pathway (model shown in **Figure 4**G).

**Figure 4.**
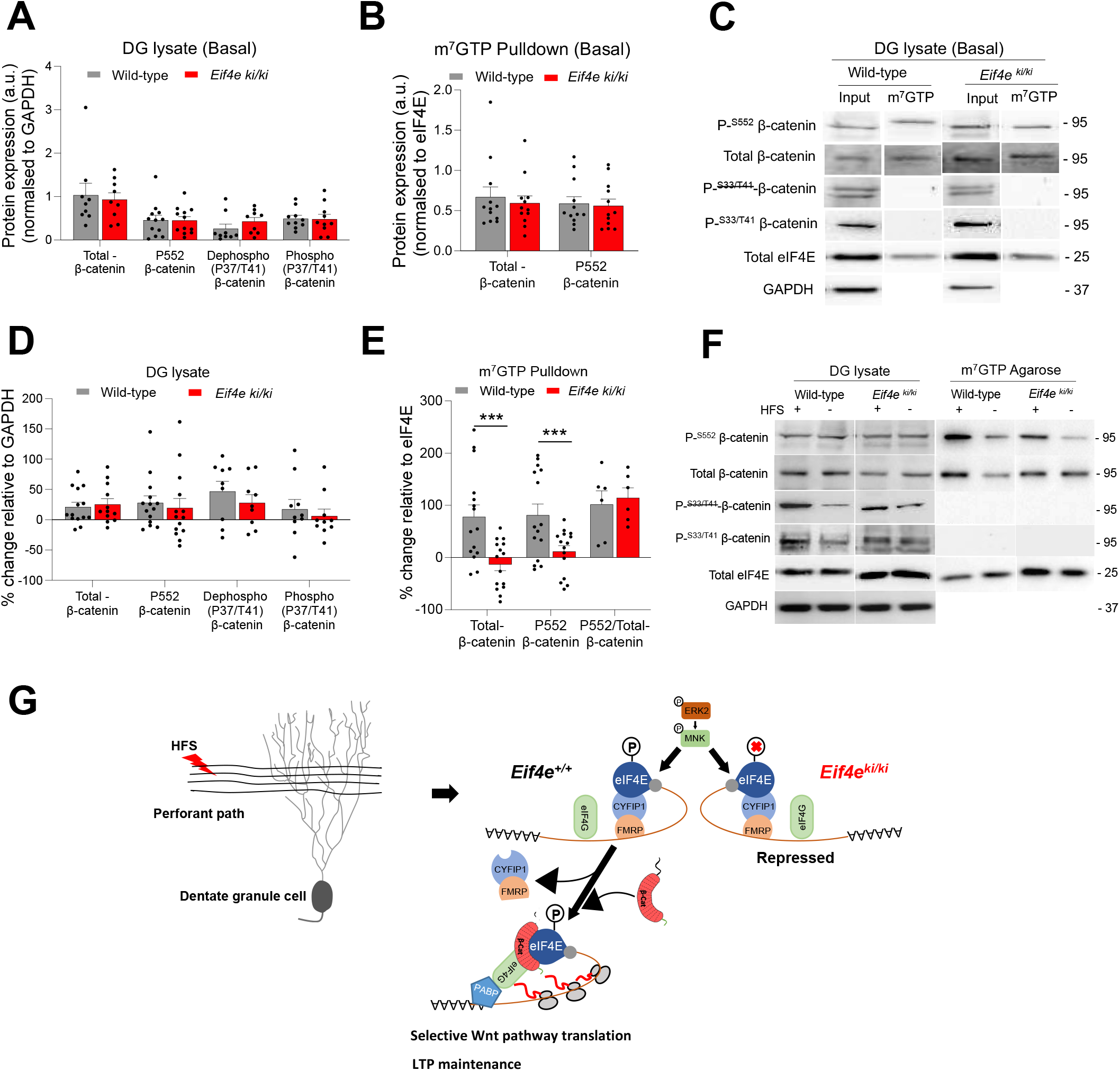
Synaptic activity-evoked eIF4E phosphorylation recruits (atypical) β-catenin to eIF4E cap complex in DG-LTP in vivo. (A) Immunoblot analysis of total (T) and phosphorylated (P) β-catenin in DG lysates from naïve mice (basal state). No significant difference between wild-type and *Eif4e*^ki/ki^ mice in expression of total β-catenin, P-552 (n=9) β-catenin Arm domain (n=12), P-37/41 β-catenin N-terminal region (n=9), or non-37/41 phosphorylated N-terminal region (n=10), normalized to GAPDH. Values are means + SEM (Multiple t-test). (B) Immunoblot analysis from m^7^GTP pulldown assays in DG lysates in naïve mice (basal state). Densitometric values for Total and P-S552-β-catenin are normalized to values of m^7^GTP-bound eIF4E. Values in *Eif4e*^ki/ki^ mice were not significantly different from wild-type. (C) Representative immunoblots for panels (A) and (B). (D) Changes in the expression of total and phosphorylated β-catenin in DG lysate following HFS. Values are expressed in percent change relative to contralateral DG control. No significant difference was observed. (E) Immunoblot analysis of m^7^GTP pulldowns from DG lysates obtained 40 min post-HFS. Total- β-cat = 0.0013, P-^S552^ β-cat = 0.0074, P-^S552^ /Total-β-cat =0.7040, (**p < 0.001, ***p < 0.0001, Multiple t-test). (Total- β-cat, n =12, P-^S552^ β-cat, n=12, P-^S552^ /Total-β-cat, n=6). (F) Representative immunoblots for panels (D) and (E). HFS = high-frequency stimulation. (+) Ipsilateral DG, (-) Contralateral DG. Mean ± SEM and Student’s t-test in Table S2. (G) Model of phospho-eIF4E-dependent translation of Wnt pathway in DG-LTP maintenance. (Left) HFS of medial perforant input to DG granule cells. (Right). In wild-type mice, synaptic activation by HFS stimulates ERK-MNK signaling and eIF4E phosphorylation on Ser209. Phosphorylation triggers discharge of eIF4E-binding proteins, both CYFIP1 and 4E-BP2, and recruitment of β-catenin (β-cat) to the eIF4E cap complex. Release of CYFIP1 together with its binding partner FMRP is depicted. The β-catenin that associates with eIF4E is atypical as its N-terminal region (indicated by dotted line) is not detected by specific antibodies. Phosphorylation of eIF4E is required for Wnt pathway translation as a class and underlies LTP maintenance. In *Eif4e*^ki/ki^ mice, ERK-MNK signaling is activated but loss of eIF4E phosphorylation prevents remodeling of the eIF4E complex, Wnt pathway translation, and LTP. See also Figure S7.

β-catenin stability and transcriptional activity are regulated by phosphorylation. We therefore asked whether the β-catenin that associates with eIF4E represents a distinct form. Immunoblotting was done using antibodies recognizing critical phosphorylation sites on β-catenin’s N-terminal intrinsically disordered region and within its central Armadillo (Arm) repeats domain (Gao et al., 2014). In the absence of Wnt, GSK3-catalyzed phosphorylation of the β-catenin N-terminus (Ser33/Thr41) promotes ubiquitination and proteasomal degradation (MacDonald et al., 2009; Nakamura et al., 2016). In the presence of Wnt, dephosphorylation of the N-terminal residues prevents degradation, allowing accumulation of β-catenin in the nucleus to regulate transcription. We probed with antibodies specifically recognizing phosphorylated (Ser33/Thr41) or non-phosphorylated N-terminal epitopes. Robust signals were detected with both antibodies in DG lysates from naïve animals, with no difference between genotypes (**Figure 4**A and 4C). Following LTP induction, enhanced expression of phosphorylated and non-phosphorylated N-terminal β-catenin were observed, but again there was no difference between knockin and wild-type (**Figure 4**D and 4F). Remarkably, immunoblots in cap-pulldown samples from naïve and HFS-treated mice of both genotypes were negative for both phosphorylated and non-phosphorylated N-terminal β-catenin (**Figure** 4C and 4F). This could mean that β-catenin in the cap complex lacks the N-terminus or is modified in a way that prohibits binding of the specific antibodies. The fact that total β-catenin is reliably detected at the expected molecular mass (95 kDa) in all cap-pulldown and lysate samples shows that β-catenin is not proteolytically cleaved.

Phosphorylation of Ser552 in the Arm domain regulates protein-protein interactions, with enhanced phosphorylation decreasing partner binding and promoting nuclear accumulation (Budnik and Salinas, 2011; Fang et al., 2007; Gao et al., 2014; Goretsky et al., 2018). β-catenin Ser552 phosphorylation was detected in both cap-pulldown and lysate samples, but again, with no difference between genotypes at baseline (**Figure 4**A-4C). Following LTP induction, phospho-Arm was increased in HFS-treated DG relative to the contralateral DG in both genotypes (**Figure 4**D and 4F). Strikingly, in cap-pulldown samples, Ser552 phosphorylated β-catenin was significantly increased 78.3 % above contralateral control in wild-type mice and this increase was abolished in *Eif4e*^ki/ki^ mice. However, normalization of phospho-to the enhanced levels of total β-catenin showed that Ser552 phosphorylation state did not change (**Figure 4**E). Thus, HFS induces a phospho-eIF4E-dependent recruitment of β-catenin to eIF4E where it maintains constitutive levels of Ser552 Arm phosphorylation.

Finally, immunoblotting in DG lysates was done to assess changes in protein expression of select, translationally upregulated Wnt pathway targets. In wild-type mice, HFS significantly increased expression of the key Wnt receptor scaffolding protein, Dvl2, relative to contralateral control, whereas no significant change was observed in *Eif4e*^ki/ki^ mice **(Figure S8**). Expression of Frizzled class receptor 4 (Fzd4) or secreted frizzled-related protein 1 (Sfrp1) was unchanged, indicating differential impacts on protein expression in early LTP maintenance at 40 min post-HFS

## DISCUSSION

This study elucidates a mechanism and function for Ser209 eIF4E phosphorylation in translational control of DG-LTP in vivo. Biochemically, eIF4E phosphorylation is required for synaptic activity-evoked remodeling of the cap-binding complex. The remodeling is bidirectional, with discharge of the translational repressors CYFIP1 and 4E-BP2, and recruitment of β-catenin. In the first translational profiling analysis of DG-LTP, we find that phospho-eIF4E is specifically required for synaptic activity-induced translation of the Wnt pathway. Functionally, ablation of phospho-eIF4E does not alter basal synaptic transmission or LTP induction but inhibits the maintenance phase of LTP. Previous work demonstrated critical roles for Wnt signaling in LTP at excitatory synapses (Chen et al., 2006; McLeod and Salinas, 2018; Tang, 2014). While β-catenin is known to mediate transcription downstream of Wnt, our results identify novel phospho-eIF4E-dependent recruitment of β-catenin to eIF4E and enhanced Wnt path translation specific to LTP maintenance.

Previous work demonstrated that acute pharmacological inhibition of MNK inhibits eIF4E phosphorylation and DG-LTP maintenance. However, MNKs have additional substrates which could impact translation and mRNA metabolism, including eIF4G, PSF (polypyrimidine tract-binding protein-associated splicing factor), and heterogenous ribonucleoprotein A1(Bramham et al., 2016; Buxadé et al., 2005; Joshi, 2014; Orton et al., 2004). Here, we show that activation of ERK-MNK signaling and LTP induction are intact in *Eif4e*^ki/ki^ mice while stable maintenance of LTP is lost. Basal perforant path-evoked synaptic transmission and eIF4E protein-protein interactions were intact, with no difference between genotypes in eIF4E associated CYFIP/FMRP, 4E-BP2, β-catenin, or eIF4G. Thus, the Ser209Ala knockin mutation does not lead to compensatory upstream changes in ERK-MNK signaling or downstream remodeling of the eIF4E complex. Rather, our results demonstrate a crucial role for phospho-eIF4E in stimulus-evoked discharge of CYFIP1 and 4E-BP2 from eIF4E and enhanced loading of eIF4G to facilitate initiation. CYFIP1 can shuttle between the eIF4E complex and a Rac-WAVE1 complex involved in actin cytoskeletal remodeling and dendritic spine plasticity (De Rubeis et al., 2013; Santini et al., 2017). If such shuttling occurs in LTP, disruption of CYFIP1 release may impact both translation and actin dynamics.

Wnts have broad and diverse functions in embryonic patterning and development, including neuronal dendrite development and synapse formation, and continue to function in activity-dependent synaptic plasticity in the adult brain (Budnik and Salinas, 2011; McLeod and Salinas, 2018; Steinhart and Angers, 2018). Neuronal activity-induced Wnt secretion and signaling through canonical β-catenin and the non-canonical planar cell polarity (PCP) and calcium pathways are implicated in LTP of excitatory synaptic transmission (Cerpa et al., 2011; Chen et al., 2006; Gogolla et al., 2009; McLeod and Salinas, 2018; Tang, 2014). Roles for Wnt signaling in trafficking of AMPA-type glutamate receptors and structural plasticity of dendritic spines have been identified (McLeod and Salinas, 2018; McLeod et al., 2018). In DG-LTP, canonical Wnt/β-catenin signaling is associated with transcription of Wnt target genes (Chen et al., 2006; Park et al., 2006). Our ribosome profiling analysis uncovered phospho-eIF4E-dependent translation of Wnt-4 and Wnt receptors and scaffolds (Dvl2, Lrp5, Fzd-2 and 4) as well as endogenous inhibitors of Wnt receptors (Sfrp1 and 2). Targeting of the Wnt pathway as a class suggests a coordinate regulation which could serve to amplify Wnt signaling in LTP.

Whether recruitment of β-catenin to eIF4E is directly involved in regulation of Wnt pathway translation remains to be determined. Our analysis of mRNA features indicates that phospho-eIF4E preferentially promotes translation of mRNAs with long, structured 5’ UTRs. However, as these features are not unique to Wnt family components, other factors must provide specificity. In the developing nervous system, β-catenin interaction with N-cadherin regulates dendritic spine plasticity (Murase et al., 2002; Yu and Malenka, 2003). N-cadherin stimulates Akt, which phosphorylates β-catenin Ser552 (Zhang et al., 2013). Phosphorylation of β-catenin Ser552 has been shown to regulate protein-protein interactions. We show that β-catenin in the eIF4E complex is Ser552 phosphorylated, but there is no change in phosphorylation state to support phosphorylation as a mechanism of recruitment. Under reducing conditions on SDS-PAGE gels, immunoblotting showed that antibodies specific for the N-terminal intrinsically disordered region reliably and clearly detect β-catenin in lysates, whereas cap-pulldown assays are blank. This suggests that β-catenin in the eIF4E complex is a unique form that has undergone a structural change or post-translational modification that prevents detection of the N-terminal epitope.

Translation control can be general, affecting translation in a global manner, or more specific, impacting only subsets of mRNAs (Dever, 2002; Gebauer and Hentze, 2004). HFS in wild-type and *Eif4e*^ki/ki^ mice increased expression of numerous IEG mRNAs and enhanced translational efficiency in gene ontology categories synaptic regulation and plasticity. Of these, the Wnt pathway was the only category of DTGs completely dependent on phospho-eIF4E. We compared our list of phospho-eIF4E regulated DTGs in DG in vivo with FMRP-target genes (Darnell et al., 2011) and MNK1 targets identified using cortical neuronal cultures from MNK1 knockout mice (**Figure S9**). While the preparations and methods are different, the main conclusion seems to be that there are few common mRNAs (less than 5%). This reinforces the importance of phospho-eIF4E in specific translation during LTP in the DG. DG-LTP consolidation has mechanistically distinct early and late phases of translation with different t targets. We concentrated on the critical, early stage of translation (40 min post-HFS) and recognize that new patterns may emerge at later time points.

In cancer models, phospho-eIF4E regulates translation of targets involved in oncogenic transformation (Furic et al., 2010; Robichaud et al., 2014). In the suprachiasmatic nucleus, phospho-eIF4E regulates translation of clock genes (*Per1* and *Per2*) involved in circadian rhythms (Cao et al., 2015) and in dorsal root ganglion phospho-eIF4E promotes translation of *bdnf* involved in hyperalgesia and nociceptive transmission (Moy et al., 2018). A recent ribosome footprinting analysis of forebrain tissue of adult *Eif4e*^ki^ mice revealed regulation of mRNAs involved in inflammation (IL-2 and TNFα) and organization of extracellular matrix (Prg2, Mmp9, Adamts16, Acan) (Amorim et al., 2018; Gkogkas et al., 2014). Behavioral analyses of *Eif4e*^ki/ki^ mice have revealed a role for phospho-eIF4E in regulation of depression-like behavior (Amorim et al., 2018). Collectively, evidence suggests that phospho-eIF4E regulates translation in a region- and stimulus-specific manner.

A previous analysis of the Schaffer collateral-CA1 pathway in hippocampal slices from *Eif4e*^ki/ki^ mice shows normal basal transmission and LTP induction as well as normal late phase LTP maintenance (recorded for 2 hours) (Amorim et al., 2018). The ability to generate stable LTP in CA1 but not DG in *Eif4e*^ki/ki^ mice could reflect differences in the balance and timing of mTORC1 and MNK-dependent translation in these circuits. LTP maintenance in CA1 requires mTORC1 signaling, which triggers removal of 4E-BP2 (Banko et al.; Briz et al., 2013; Stoica et al., 2011; Tang et al., 2002). Stable CA1-LTP induced by HFS or BDNF application is blocked by the mTORC1 inhibitor rapamycin (Cammalleri et al., 2003; Tang et al., 2002). In DG-LTP, inhibition of mTORC1 signaling by rapamycin does not affect LTP induction or maintenance (Panja et al., 2009, 2014). It is therefore possible that ablation of phospho-eIF4E is compensated by mTORC1 signaling in region CA1, whereas MNK regulation predominates in DG-LTP and is not developmentally compensated in *Eif4e*^ki/ki^ mice.

A recent study showed that inhibition of eukaryotic elongation factor 2 (eEF2) phosphorylation by conditional deletion of eEF2 kinase dramatically increases neurogenesis in the adult DG, enhances DG-dependent cognitive functions and decreases depression-like behavior (Taha et al., 2020). The regulation of neurogenesis is specifically linked to proteostasis of mature DG granule cells, with increased expression of neurogenesis-related proteins (decorin, vimentin). The phenotype in *Eif4e*^ki/ki^ mice is very different, with normal DG neurogenesis and hippocampal-dependent memory function, while depression-like behavior is increased (Aguilar-valles et al., 2018; Amorim et al., 2018). Conceivably, eEF2 has a primary function in supporting neurogenesis-related translation while phosho-eIF4E supports synaptic plasticity of pre-existing inputs to mature granule cells. These different forms of translational control may cooperate in DG functions such as regulation of depression-like behavior.

## Methods

### STAR METHODS

#### Animals

*Eif4e^S209A^* mice previously described (Gkogkas et al., 2014) were used. In vivo electrophysiological experiments were carried out on 90 male homozygous *Eif4e*^S209A/S209A^ mice (*Eif4e*^ki/ki^), heterozygous *Eif4e*^ki/+^mice, and wild-type C57BL/6 mice (*Eif4e*^+/+^) (Taconic Europe, Ejby, Denmark), weighing 25-30 g. Mice were bred and housed in their home cages. Room temperature (22°C±1°C) and relative humidity (46±5%) was maintained. Mice had free access to water and autoclaved standard rodent diet (SDS, England; RMI-E) and were maintained on a 12 h light/dark cycle. This research is approved by Norwegian National Research Ethics Committee in compliance with EU Directive 2010/63/EU, ARRIVE guidelines. Persons involved in the animal experiments have approved Federation of Laboratory and Animal Science Associations (FELASA) C course certificates and training.

#### Antibodies used

Antibodies used for immunoblotting are listed in **Table S1**.

#### *In vivo* electrophysiology in mice

Electrophysiology methods are as described (Panja et al., 2014) with minor modifications. Adult mice (12-weeks old) were anesthetized with urethane (injected i.p. 1.2 g/kg), which was supplemented throughout surgery and recording as required. Mice were placed in a stereotaxic frame and body temperature was maintained at 37°C. In one hemisphere only, A bipolar stimulation electrode (NE-200, 0.5 mm tip separation, Rhodes Medical Instruments, Wood hills, CA) was positioned for unilateral stimulation of the perforant path (3.8 mm posterior to bregma, 2.5 mm lateral to midline, and 1.6 mm depth from the brain surface) while an insulated tungsten recording electrode (0.075 mm; A-M Systems) was positioned in the DG hilar region (2 mm posterior to bregma, 1.5 mm lateral to the midline, and 1.5 – 1.7 mm depth from the brain surface). The recording electrode was lowered into the brain in 0.1 mm increments while monitoring the laminar profile of the response waveform evoked by a 400 μA test-pulse stimulus. To generate input/output (I/O) curves, 7 stimulus intensities ranging from 80 μA to 400 μA were applied in randomized sequence. After generating an I/O curve, a stable 20 min baseline of evoked potentials was recorded using test-pulses of 0.1 ms pulsewidth applied at 0.033 Hz. The test-pulses intensity produced a population spike of 30% of maximum. The high-frequency stimulation (HFS) protocol consisted of four trains of stimuli applied with an interval of 10 sec; each train had 15 pulses at 200 Hz (pulse-width 0.1 ms). The stimulus intensity for HFS was twice that used for baseline recordings. After HFS, test-pulse evoked responses were recorded for 40- and 180-min. After recordings were completed, the electrodes were removed, the animal was sacrificed, and the dentate gyri were microdissected and immediately frozen on dry-ice for later use. The maximal slope of the initial rising phase of the fEPSP and population spike amplitude were measured, changes post-HFS were expressed in percent of baseline. In the electrophysiological experiments for ribosome profiling, mice received the standard LTP protocol consisting of low-frequency test-pulse stimulation (LFS) and HFS. DG tissue was collected at 40 min post-HFS. To control for effects of test-pulse stimulation, a control group received LFS only for 40 min.

#### Tissue dissection and sample preparation

At the end of electrophysiological recording, ipsilateral and contralateral dentate gyri were rapidly dissected on ice and homogenized in buffer containing 50 mM Tris, 100 mM NaCl, 1 mM EDTA, NP-40 0.5%, 1 mM dithiothreitol, 1 mM Na_3_VO_4_, 50 mM NaF, and 1× protease inhibitor cocktail from Roche #11836170001. Homogenization was performed manually with 10–12 gentle strokes in a tissue grinder and the homogenate was centrifuged for 10 min at 14000 x g at 4°C. Protein concentration was measured using the BCA protein assay (Pierce, #23227). Homogenates were stored at −80°C until use.

#### m^7^GTP pulldown assays

m^7^GTP pulldown assays have been described in detailed elsewhere (Amorim et al., 2018; Panja et al., 2014). In brief, 250-300 μg of protein lysate together with 30 μl of 7-methyl GTP-agarose beads (Jena bioscience #AC-141) were incubated for 90 min at 4°C. Beads were washed three times with m^7^GTP lysis buffer and bound proteins were separated to an SDS-PAGE (10% gels or 4-15% gradient gels). Immunoblotting was carried out as described below.

#### SDS–PAGE and immunoblotting

Samples from m^7^GTP pull-down assays and lysates were boiled at 70°C for 10 min in Laemmli sample buffer (Bio-Rad) and resolved in 10% or 4-15% gradient SDS-PAGE gels. Proteins were transferred to nitrocellulose membranes (Biorad, # 162-0112) which were then blocked with 5% BSA, probed with antibodies and developed using chemiluminescence reagents (Biorad, #1705061). The blots were scanned using Gel DOC XRS+ (BIO RAD) and densitometric analyses were performed with Image J software (NIH, Bethesda, MD). Blots treated with phospho-specific antibody stripped with 100 mM 2-mercaptoethanol, 2% SDS and 62.5 mM Tris-HCl, pH 6.8 at 50° C for 45 min, washed, blocked and re-probed with antibody recognizing total protein. Densitometric values expressed per unit of protein applied to the gel lane. Total proteins normalized to loading control. Values from the HFS-treated DG were expressed in percent change of values from contralateral control DG. Significant differences between the treated and non-treated animal were determined using one-way ANOVA or Student’s *t*-test for dependent samples. The p-value for significance was 0.05.

#### Ribosome Profiling (RP) & Bioinformatics Analysis

Tissue was processed using the TruSeq Ribo Profile (Mammalian) Kit (Illumina), according to the manufacturer’s instructions. Briefly, tissue was homogenised in Mammalian Polysome Buffer (Illumina) supplemented with DNAse I (10 U/mL), 1% Triton X-100, 1% NP-40, 1 mM DTT and cycloheximide (100 μg/mL). Half of the lysate was used for mRNA extraction (total mRNA) while the remaining fraction was digested with TruSeq Ribo Profile Nuclease so that only the mRNA fragments protected by ribosomes were recovered (footprints). Both samples (footprints and total mRNA) went through a ribosomal RNA removal step using the Ribo-Zero Gold (Human/Mouse/Rat) Kit (Illumina). The footprint samples were then purified on a 15% TBE-Urea polyacrylamide gel (ThermoFisher Scientific) to select for fragments of 28-30 nucleotides. The total mRNA samples were heat-fragmented, according to the TruSeq Ribo Profile protocol, to yield small RNA fragments. Footprints and total mRNA fragments were used to prepare small RNA libraries, using the TruSeq Ribo Profile Kit, and were sequenced on an Illumina HiSeq 2500 System, at the Edinburgh Genomics facilities. Bioinformatics analysis was performed as previously described (Amorim et al., 2018). Translational Efficiency (TE) was calculated as the ratio between RPKM of footprints and RPKM of total mRNA for each gene. Data was filtered to include only Differentially Translated Genes (DTGs) that meet the following criteria: FDR<0.05, p-value <0.05, and - 1<Log2(TE)>0.585.

#### Gene Ontology and Pathway Analysis

Gene Ontology (GO) and Pathway Analysis were performed using the online tool DAVID (Database for Annotation, Visualization and Integrated Discovery; (Huang et al., 2009) version 6.8) and the Ingenuity Pathway Analysis Software (IPA; Qiagen; version 42012434), respectively. Differentially translated genes were submitted to IPA and subjected to Core Analysis with analysis parameters set to include Direct and Indirect Interactions and Experimentally Observed data only. For further analysis of relevant Canonical Pathways, a Molecular Activity Predictor (MAP) analysis was applied based on the differentially regulated genes belonging to each individual pathway. For GO analysis, filtered gene lists split to highlight genes differentially upregulated or downregulated in each dataset were individually submitted to DAVID and GO annotation gathered for Biological Function, Molecular Function and Cellular Component.

#### Statistical analysis

Group values are reported as mean ± SEM. Statistical comparisons were calculated with the Two-way repeated measures ANOVA with Bonferroni multiple-comparison test or multiple T-test with Holm-Sidak method using GraphPad Prism 8.02. Significance level was set *a priori* at p < 0.05. Statistical analysis data is shown in Table S2.

## ACKNOWLEDGMENTS

eIF4E^Ser209Ala^ mice were generously provided by Prof. Nahum Sonenberg, McGill University, Montreal, Canada. Work in the C.R.B lab was supported by Research Council of Norway (RCN) grants 249951 and 226026, and the EU Joint Programme - Neurodegenerative Disease Research (JPND) project, CircProt (project 30122) in which funding to C.R.B. was provided by RCN. S.P. and T.F.M. received fellowships from the University of Bergen, Medical Faculty. Work in the C.G.G. lab was supported by the Hellenic Foundation for Research and Innovation (H.F.R.I.) under the “2nd Call for H.F.R.I. Research Projects to support Faculty Members & Researchers” (Project Number: 2556).

## AUTHOR CONTRIBUTIONS

S.P., T.F.M. and S.A. conducted the in vivo electrophysiology experiments. S.P. and T.F.M. conducted the biochemical experiments. K.C., K.S., and I.S. conducted the ribosome profiling. S.P., K.C. and K.S. analyzed data and performed bioinformatic analysis. S.P., C.G.G, and C.R.B conceived the study and wrote the paper.

## SUPPLEMENTAL INFORMATION

### Supplemental Figure Legends

**Figure S1. Input-output curves of medial perforant path-DG evoked responses show normal basal synaptic transmission and excitability in *Eif4e*^ki/ki^ mice.**

(A) Input-output relationship of the fEPSP slope in *Eif4e*^ki/ki^ and wild-type littermates. Current intensities ranged from 80-300 μA. No significant difference between groups (n = 4, p = 0.9974; Two-way ANOVA).

(B) Population spike (Spike) amplitude (p = 0.5977).

(C) EPSP-Spike plot. p= 0.7631). Values are mean ± SEM.

**Figure S2. Population spike LTP in *Eif4e*^ki/ki^ mice.**

(A) Time-course plots of medial perforant path-DG evoked population spike amplitude recorded before and after high-frequency stimulation (HFS, indicated by arrow) in homozygous *Eif4e*^S209A^ knockin mice (*Eif4e*^ki/ki^; n=7), heterozygous mice (*Eif4e*^+/ki^; n=8) and wild-type littermates (*Eif4e^+/+^* mice; n=10). Values are mean (± SEM) expressed in percent of baseline.

(B) Bar graphs of mean changes in population spike amplitude recorded between 0-10 min, 30-40 min, and 170-180 min post-HFS. There was no significant difference between genotypes at these time points. Student’s t-test.

Figure 1C shows representative field potentials.

**Figure S3. Basal and post-HFS expression of translation factors in DG lysates.**

(A) Immunoblot analysis eIF4G, eIF4E, 4E-BP2, CYFIP1, FMRP and Arc in DG lysates from naïve mice (basal state). Expression is normalized GAPDH. No significant difference between wild-type and *Eif4e*^ki/ki^ mice. Values are means + SEM.

(B) Immunoblot analysis of eIF4G (n=12, 0.02234), eIF4E (n=16, 0.7712), 4E-BP2 (n=11, 0.0756), CYFIP1 (n=14, 0.00007), FMRP (n=8, 0.0072) and Arc (n=17, 0.018) in HFS-treated DG. Values are mean (± SEM) expressed as percent change relative to contralateral DG. Significant differences between genotypes are indicated. (*p < 0.05, **p < 0.001, ***p < 0.0001, Multiple t-test).

(C) Representative immunoblots for panel (B).

**Figure S4. Quality Control (QC) of ribosome profiling experiment.**

(A) Graphical depiction of groups presented in this figure; LFS: low frequency stimulation, HFS: high frequency stimulation; in *Eif4e*^ki/ki^ and *Eif4e^+/+^* (wild-type) mice ribosome profiling experiment in Fig. 3.

(B) (Top panel) Size of footprints and total mRNA libraries analysed; frequency of different size fragments (length of mapped reads; bp). (Bottom) Fraction of reads within start codon window for frames 1, 2 and 3.

(C) Periodicity of reads (frequency) around the start and stop codons (nt from) for the ribosome profiling experiment (footprints & total mRNA) for the experimental groups shown (see legend for A.).

**Figure S5. Comparison of transcription, translation and gene ontology in HFS vs contralateral control and test-pulse LFS control in *Eif4e^+/+^* (wild-type) mice.**

**A.** Scatter plot of TE or RPKM comparisons for the indicated groups: low-frequency test-pulse stimulation (LFS); standard LTP protocol of test-pulse stimulation and high-frequency stimulation (HFS); in *Eif4e*^ki/ki^ and *Eif4e*^+/+^ (wild-type) mice. HFS-treated ipsilateral DG is compared to contralateral, non-stimulated DG and to ipsilateral DG of mice given LFS only.

**B.** Gene Ontology analysis using IPA canonical pathways for DTGs and DEG. Top 3 groups are shown.

**Figure S6. Gene ontology analysis for upregulated DTGs in *Eif4e*^ki/ki^ mice from ribosome profiling experiment in Fig. 3**

Gene ontology analysis of downregulated genes in *Eif4e^+/+^* mice (57 genes) and *Eif4e*^ki/ki^ mice (91 genes); plots for biological process (A) cellular component (B) and molecular function

(C) with number of genes in each category with p-values. (D) KEGG pathway analysis for upregulated genes. Mean ± SEM and Student’s t-test in **Table S2**.

**Figure S7. 5’-UTR sequence analysis using UTRscan.**

(A) (TOP) Summary table depicting differences in the variables (length, %GC and Gibbs Free energy) shown in bar graphs. (BOTTOM) Length (nt; nucleotides), %GC content and Gibbs free energy (kcal/mol) are shown for upregulated and downregulated DTGs from ribosome profiling experiment (WT 40 min HFS and KI 40 min HFS).

(B) (TOP) Summary table depicting differences in motifs (uORF, TOP, IRES, PG4) shown in bar graphs. (BOTTOM) Bar graphs for UTRscan motif analysis in for upregulated and downregulated DTGs from ribosome profiling experiment (WT 40 min HFS and KI 40 min HFS). Data are shown as mean ± SEM. For A; Mann-Whitney test ***p<0.001, *p<0.05.

**Figure S8. Immunoblot analysis validates phospho-eIF4E-dependent expression of Wnt pathway receptor, dishevelled 2 (Dvl2), following LTP induction in the dentate gyrus.**

A) Immunoblot analysis of Dvl2, Fzd4, and Sfrp1 in DG lysates from naïve mice (basal state). Densitometric values are normalized to GAPDH. Values are means + SEM. No significant difference between wild-type and *Eif4e*^ki/ki^ mice.

(B) Changes in the expression of Dvl2, Fzd4, and Sfrp1 in DG lysate following HFS. Values are expressed in percent change relative to contralateral DG control. Gives stats for genotype comparison.

(C) Representative immunoblots for panel (B). HFS = high-frequency stimulation. (+) Ipsilateral DG, (-) Contralateral DG.

**Figure S9. Comparison of Patil et al. data with previous studies.**

Venn diagrams showing comparison of ribosome profiling DTGs at baseline or 40 min post-HFS (*Eif4e*^+/+^ vs *Eif4e*^ki/ki^) from the present study with (**A)** 842 FMRP HITS-CLIP targets (Darnell et al., 2011) and (**B)** 91Mnk1 targets identified by BONCAT and SILAC labeling in BDNF-treated cultured cortical neurons (Genheden et al., 2015).

### Supplementary Tables

**Table S1. Antibodies and reagents**

**Table S2. Statistics.**

**Table S3. Differential expression and translation of genes in HFS-treated wild-type mice: comparison with contralateral and LFS-treated control groups.**

**Table S4. Gene Ontology (GO) and Pathway Analysis.**

**Table S5**. **Differential expression and translation of genes in HFS-treated wild-type *Eif4e^+/+^* mice and *Eif4e*^ki/ki^ mice.**

**Table S6. List of differentially translated mRNAs in GO categories canonical Wnt signaling, actin cytoskeletal regulation, and ribosomal proteins.**

## REFERENCES

Aguilar-valles, A., Haji, N., Gregorio, D. De, Matta-camacho, E., Eslamizade, M.J., Popic, J., Sharma, V., Cao, R., Rummel, C., Tanti, A., et al. (2018). Translational control of depression-like behavior via phosphorylation of eukaryotic translation initiation factor 4E. Nat. Commun.

Amorim, I.S., Kedia, S., Kouloulia, S., Simbriger, K., Gantois, I., Jafarnejad, S.M., Li, Y., Kampaite, A., Pooters, T., Romanò, N., et al. (2018). Loss of eIF4E phosphorylation engenders depression-like behaviors via selective mRNA translation. J. Neurosci. 38, 2118–2133.

Banko, J.L., Poulin, F., Hou, L., DeMaria, C.T., Sonenberg, N., and Klann, E. The translation repressor 4E-BP2 is critical for eIF4F complex formation, synaptic plasticity, and memory in the hippocampus. J. Neurosci. 25, 9581–9590.

Beggs, J.E., Tian, S., Jones, G.G., Xie, J., Iadevaia, V., Jenei, V., Thomas, G., and Proud, C.G. (2015). The MAP kinase-interacting kinases regulate cell migration, vimentin expression and eIF4E/CYFIP1 binding. Biochem. J. 467, 63–76.

Bramham, C.R., Jensen, K.B., and Proud, C.G. (2016). Tuning specific translation in cancer metastasis and synaptic memory: control at the MNK – eIF4E axis. Trends Biochem. Sci. 41, 847–858.

Briz, V., Hsu, Y.-T., Li, Y., Lee, E., Bi, X., and Baudry, M. (2013). Calpain-2-Mediated PTEN Degradation Contributes to BDNF-Induced Stimulation of Dendritic Protein Synthesis. J. Neurosci 33, 4317–4328.

Budnik, V., and Salinas, P.C. (2011). Wnt signaling during synaptic development and plasticity. Curr. Opin. Neurobiol. 21, 151–159.

Buxadé, M., Parra, J.L., Rousseau, S., Shpiro, N., Marquez, R., Morrice, N., Bain, J., Espel, E., and Proud, C.G. (2005). The Mnks are novel components in the control of TNFα biosynthesis and phosphorylate and regulate hnRNP A1. Immunity 23, 177–189.

Cammalleri, M., Lütjens, R., Berton, F., King, A.R., Simpson, C., Francesconi, W., and Sanna, P.P. (2003). Time-restricted role for dendritic activation of the mTOR-p70S6K pathway in the induction of late-phase long-term potentiation in the CA1. Proc. Natl. Acad. Sci. U. S. A. 100, 14368–14373.

Cao, R., Gkogkas, C.G., de Zavalia, N., Blum, I.D., Yanagiya, A., Tsukumo, Y., Xu, H., Lee, C., Storch, K.-F., Liu, A.C., et al. (2015). Light-regulated translational control of circadian behavior by eIF4E phosphorylation. Nat. Neurosci. 18.

Cerpa, W., Gambrill, A., Inestrosa, N.C., and Barria, A. (2011). Regulation of NMDA-receptor synaptic transmission by Wnt signaling. J. Neurosci. 31, 9466–9471.

Chalkiadaki, K., Kouloulia, S., Bramham, C.R., and Gkogkas, C.G. (2021). Regulation of Protein Synthesis by eIF4E in the Brain. In The Oxford Handbook of Neuronal Protein Synthesis, W.S. Sossin, ed. (Oxford University Press), pp. 2–22.

Chen, J., Chang, S.P., and Tang, S.J. (2006). Activity-dependent synaptic Wnt release regulates hippocampal long term potentiation. J. Biol. Chem. 281, 11910–11916.

Chen, P.B., Kawaguchi, R., Blum, C., Achiro, J.M., Coppola, G., O’Dell, T.J., and Martin, K. C. (2017). Mapping gene expression in excitatory neurons during hippocampal late-phase long-term potentiation. Front. Mol. Neurosci. 10, 1–16.

Costa-Mattioli, M., Sossin, W.S., Klann, E., and Sonenberg, N. (2009). Translational control of long-lasting synaptic plasticity and memory. Neuron 61, 10–26.

Darnell, J.C., Van Driesche, S.J., Zhang, C., Hung, K.Y., Mele, A., Fraser, C.E., Stone, E.F., Chen, C., Fak, J.J., Chi, S.W., et al. (2011). FMRP stalls ribosomal translocation on mRNAs linked to synaptic function and autism. Cell. 146, 247–261.

Dever, T.E. (2002). Gene-specific regulation by general translation factors. Cell 108, 545–556.

Ehyai, S., Miyake, T., Vinayak, J., McDermott, J.C., Ehyai, S., Bayfield, M.A., and Williams, D. (2018). FMRP recruitment of β□catenin to the translation pre□initiation complex represses translation. EMBO Rep. 19, e45536.

Fang, D., Hawke, D., Zheng, Y., Xia, Y., Meisenhelder, J., Nika, H., Mills, G.B., Kobayashi, R., Hunter, T., and Lu, Z. (2007). Phosphorylation of ß-catenin by AKT promotes β-catenin transcriptional activity. J. Biol. Chem. 282, 11221–11229.

Furic, L., Rong, L., Larsson, O., Koumakpayi, I.H., Yoshida, K., Brueschke, A., Petroulakis, E., Robichaud, N., Pollak, M., Gaboury, L.A., et al. (2010). eIF4E phosphorylation promotes tumorigenesis and is associated with prostate cancer progression. Proc Natl Acad Sci USA 107, 14134–14139.

Gal-Ben-Ari, S., Kenney, J.W., Ounalla-Saad, H., Taha, E., David, O., Levitan, D., Gildish, I., Panja, D., Pai, B., Wibrand, K., et al. (2012). Consolidation and translation regulation. Learn. Mem. 19, 410–422.

Gao, C., Xiao, G., and Hu, J. (2014). Regulation of Wnt/βcatenin signaling by post-translational modifications. Cell Biosci. 4, 1–20.

Gebauer, F., and Hentze, M.W. (2004). Molecular mechanisms of translational control. Nat. Rev. Mol. Cell Biol. 5, 827–835.

Gelinas, J.N., Banko, J.L., Hou, L., Sonenberg, N., Weeber, E.J., Klann, E., and Nguyen, P. V (2007). ERK and mTOR signaling couple beta-adrenergic receptors to translation initiation machinery to gate induction of protein synthesis-dependent long-term potentiation. J. Biol. Chem. 282, 27527–27535.

Genheden, M., Kenney, J.W., Johnston, X.H.E., Manousopoulou, A., Garbis, S.D., and Proud, X.C.G. (2015). BDNF stimulation of protein synthesis in cortical neurons requires the MAP kinase-interacting kinase MNK1. J Neurosci 35, 972–984.

Gingras, A.C., Raught, B., and Sonenberg, N. (2001). Control of translation by the target of rapamycin proteins. Prog.Mol.Subcell.Biol. 27, 143–174.

Gkogkas, C.G., Khoutorsky, A., Cao, R., Jafarnejad, S.M., Prager-Khoutorsky, M., Giannakas, N., Kaminari, A., Fragkouli, A., Nader, K., Price, T.J., et al. (2014). Pharmacogenetic inhibition of eIF4E-dependent Mmp9 mRNA translation reverses fragile X syndrome-like phenotypes. Cell Rep. 9, 1–14.

Gogolla, N., Galimberti, I., Deguchi, Y., and Caroni, P. (2009). Wnt Signaling Mediates Experience-Related Regulation of Synapse Numbers and Mossy Fiber Connectivities in the Adult Hippocampus. Neuron 62, 510–525.

Goretsky, T., Bradford, E.M., Ye, Q., Lamping, O.F., Vanagunas, T., Moyer, M.P., Keller, P.C., Sinh, P., Llovet, J.M., Gao, T., et al. (2018). Beta-catenin cleavage enhances transcriptional activation. Sci. Rep. 8, 1–15.

Havik, B., Rokke, H., Dagyte, G., Stavrum, A.K., Bramham, C.R., and Steen, V.M. (2007). Synaptic activity-induced global gene expression patterns in the dentate gyrus of adult behaving rats: Induction of immunity-linked genes. Neuroscience..

Hien, A., Molinaro, G., Liu, B., Huber, K.M., and Richter, J.D. (2020). Ribosome profiling in mouse hippocampus: plasticity-induced regulation and bidirectional control by TSC2 and FMRP. Mol. Autism 11, 1–18.

Huang, D.W., Sherman, B.T., and Lempicki, R.A. (2009). Systematic and integrative analysis of large gene lists using DAVID bioinformatics resources. Nat. Protoc. 4, 44–57.

Joshi, S. (2014). Mnk kinase pathway: Cellular functions and biological outcomes. World J. Biol. Chem. 5, 321.

Josselyn, S.A., and Tonegawa, S. (2020). Memory engrams: Recalling the past and imagining the future. Science (80-.). 367, eaaw4325.

Kelleher, R.J., Govindarajan, A., and Tonegawa, S. (2004). Translational regulatory mechanisms in persistent forms of synaptic plasticity. Neuron 44, 59–73.

Klann, E., and Dever, T.E. (2004). Biochemical mechanisms for translational regulation in synaptic plasticity. Nat. Rev. Neurosci. 5, 931–942.

Kong, J., and Lasko, P. (2012). Translational control in cellular and developmental processes. Nat. Rev. Genet. 13, 383–394.

Maag, J.L. V., Panja, D., Sporild, I., Patil, S., Kaczorowski, D.C., Bramham, C.R., Dinger, M.E., and Wibrand, K. (2015). Dynamic expression of long noncoding RNAs and repeat elements. Front. Neurosci. 9, 1–16.

MacDonald, B.T., Tamai, K., and He, X. (2009). Wnt/ß-Catenin Signaling: Components, Mechanisms, and Diseases. Dev. Cell 17, 9–26.

Matthies, H., Frey, U., Reymann, K., Krug, M., Jork, R., and Schroeder, H. (1990). Different mechanisms and multiple stages of LTP. Adv.Exp.Med.Biol. 268, 359–368.

McLeod, F., and Salinas, P.C. (2018). Wnt proteins as modulators of synaptic plasticity. Curr. Opin. Neurobiol. 53, 90–95.

McLeod, F., Bossio, A., Marzo, A., Ciani, L., Sibilla, S., Hannan, S., Wilson, G.A., Palomer, E., Smart, T.G., Gibb, A., et al. (2018). Wnt Signaling Mediates LTP-Dependent Spine Plasticity and AMPAR Localization through Frizzled-7 Receptors. Cell Rep. 23, 1060–1071.

Messaoudi, E., Kanhema, T., Soulé, J., Tiron, A., Dagyte, G., da Silva, B., and Bramham, C.R. (2007). Sustained Arc/Arg3.1 synthesis controls long-term potentiation consolidation through regulation of local actin polymerization in the dentate gyrus in vivo. J. Neurosci 27, 10445–10455.

Moy, J.K., Khoutorsky, A., Asiedu, M.N., Dussor, G., and Price, T.J. (2018). EIF4E phosphorylation influences Bdnf mRNA translation in mouse dorsal root ganglion neurons. Front. Cell. Neurosci. 12, 1–10.

Murase, S., Mosser, E., and Schuman, E.M. (2002). Depolarization drives beta-Catenin into neuronal spines promoting changes in synaptic structure and function. Neuron 35, 91–105.

Nakamura, Y., De Paiva Alves, E., Veenstra, G.J.C., and Hoppler, S. (2016). Tissue-and stage-specific Wnt target gene expression is controlled subsequent to β-catenin recruitment to cis-regulatory modules. Dev. 143, 1914–1925.

Nguyen, P. V, and Kandel, E.R. (1996). A macromolecular synthesis-dependent late phase of long-term potentiation requiring cAMP in the medial perforant pathway of rat hippocampal slices. J. Neurosci. 16, 3189–3198.

Nguyen, T.M., Kabotyanski, E.B., Dou, Y., Reineke, L.C., Zhang, P., Zhang, X.H.F., Malovannaya, A., Jung, S.Y., Mo, Q., Roarty, K.P., et al. (2018). FGFR1-activated translation of WNT pathway components with structured 50 utrs is vulnerable to inhibition of EIF4A-dependent translation initiation. Cancer Res. 78, 4229–4240.

Nusse, R., and Clevers, H. (2017). Wnt/ß-Catenin Signaling, Disease, and Emerging Therapeutic Modalities. Cell 169, 985–999.

Orton, K.C., Ling, J., Waskiewicz, A.J., Cooper, J.A., Merrick, W.C., Korneeva, N.L., Rhoads, R.E., Sonenberg, N., and Traugh, J.A. (2004). Phosphorylation of Mnk1 by caspase-activated Pak2/γ-PAK inhibits phosphorylation and interaction of eIF4G with Mnk. J. Biol. Chem. 279, 38649–38657.

Panja, D., and Bramham, C.R. (2014). BDNF mechanisms in late LTP formation: A synthesis and breakdown. Neuropharmacology 76 Pt C, 664–676.

Panja, D., Dagyte, G., Bidinosti, M., Wibrand, K., Kristiansen, A.-M., Sonenberg, N., and Bramham, C.R. (2009). Novel translational control in Arc-dependent long term potentiation consolidation in vivo. J. Biol. Chem. 284, 31498–31511.

Panja, D., Kenney, J.W., D’Andrea, L., Zalfa, F., Vedeler, A., Wibrand, K., Fukunaga, R., Bagni, C., Proud, C.G., and Bramham, C.R. (2014). Two-stage translational control of dentate gyrus LTP consolidation is mediated by sustained BDNF-TrkB signaling to MNK. Cell Rep. 9, 1430–1445.

Park, C.S., Gong, R., Stuart, J., and Tang, S.J. (2006). Molecular network and chromosomal clustering of genes involved in synaptic plasticity in the hippocampus. J.Biol.Chem. 281, 30195–30211.

Proud, C.G. (2009). mTORC1 signalling and mRNA translation. Biochem.Soc.Trans. 37, 227–231.

Robichaud, N., Del Rincon, S. V, Huor, B., Alain, T., Petruccelli, L.A., Hearnden, J., Goncalves, C., Grotegut, S., Spruck, C.H., Furic, L., et al. (2014). Phosphorylation of eIF4E promotes EMT and metastasis via translational control of SNAIL and MMP-3. Oncogene 34, 1–11.

De Rubeis, S., Pasciuto, E., Li, K.W., Fernández, E., Di Marino, D., Buzzi, A., Ostroff, L.E., Klann, E., Zwartkruis, F.J.T., Komiyama, N.H., et al. (2013). CYFIP1 Coordinates mRNA translation and cytoskeleton remodeling to ensure proper dendritic spine formation. Neuron 79, 1169–1182.

Ryan, M.M., Mason-Parker, S.E., Tate, W.P., Abraham, W.C., and Williams, J.M. (2011). Rapidly induced gene networks following induction of long-term potentiation at perforant path synapses in vivo. Hippocampus 21, 541–553.

Santini, E., Huynh, T.N., Longo, F., Koo, S.Y., Mojica, E., D’Andrea, L., Bagni, C., and Klann, E. (2017). Reducing eIF4E-eIF4G interactions restores the balance between protein synthesis and actin dynamics in fragile X syndrome model mice. Sci. Signal. 10, 41–46.

Sonenberg, N., and Hinnebusch, A.G. (2009). Regulation of translation initiation in eukaryotes: mechanisms and biological targets. Cell 136, 731–745.

Steinhart, Z., and Angers, S. (2018). Wnt signaling in development and tissue homeostasis. Development 145, 1–8.

Stoica, L., Zhu, P.J., Huang, W., Zhou, H., Kozma, S.C., and Costa-Mattioli, M. (2011). Selective pharmacogenetic inhibition of mammalian target of Rapamycin complex I (mTORC1) blocks long-term synaptic plasticity and memory storage. Proc. Natl. Acad. Sci. U. S. A. 108, 3791–3796.

Taha, E., Patil, S., Barrera, I., Proud, C.G., Bramham, C.R., and Rosenblum, K. (2020). eEF2/eEF2K pathway in the mature dentate gyrus determines neurogenesis level and cognition. Curr. Biol. 30, 1–15.

Tang, S.-J. (2014). Synaptic activity-regulated Wnt signaling in synaptic plasticity, glial function and chronic pain. CNS Neurol. Disord. - Drug Targets 13, 737–744.

Tang, S.J., Reis, G., Kang, H., Gingras, A.C., Sonenberg, N., and Schuman, E.M. (2002). A rapamycin-sensitive signaling pathway contributes to long-term synaptic plasticity in the hippocampus. Proc.Natl.Acad.Sci.U.S.A 99, 467–472.

Wang, X., Flynn, A., Waskiewicz, A.J., Webb, B., Vries, R.G., Baines, I.A., Cooper, J.A., and Proud, C.G. (1998). The Phosphorylation of eukaryotic initiation factor eIF4E in response to phorbol esters, cell stresses, and cytokines is mediated by distinct MAP kinase pathways. J. Biol. Chem. 273, 9373–9377.

Waskiewicz, A.J., Flynn, A., Proud, C.G., and Cooper, J.A. (1997). Mitogen-activated protein kinases activate the serine/threonine kinases Mnk1 and Mnk2. EMBO J. 16, 1909–1920.

Wibrand, K., Messaoudi, E., Håvik, B., Steenslid, V., Løvlie, R., Steen, V.M., and Bramham, C.R. (2006). Identification of genes co-upregulated with Arc during BDNF-induced long-term potentiation in adult rat dentate gyrus in vivo. Eur. J. Neurosci. 23, 1501–1511.

Yu, X., and Malenka, R.C. (2003). ß-Catenin is critical for dendritic morphogenesis. Nat. Neurosci. 6, 1169–1177.

Zhang, J., Shemezis, J.R., McQuinn, E.R., Wang, J., Sverdlov, M., and Chenn, A. (2013). AKT activation by N-cadherin regulates beta-catenin signaling and neuronal differentiation during cortical development. Neural Dev. 8, 1–16.

