## Supplementary figures and images for "eIF4E phosphorylation recruits β-catenin to mRNA cap and selectively promotes Wnt pathway translation in dentate gyrus LTP maintenance in vivo"

### Supp Fig S1

#### Supplemental Figure S1

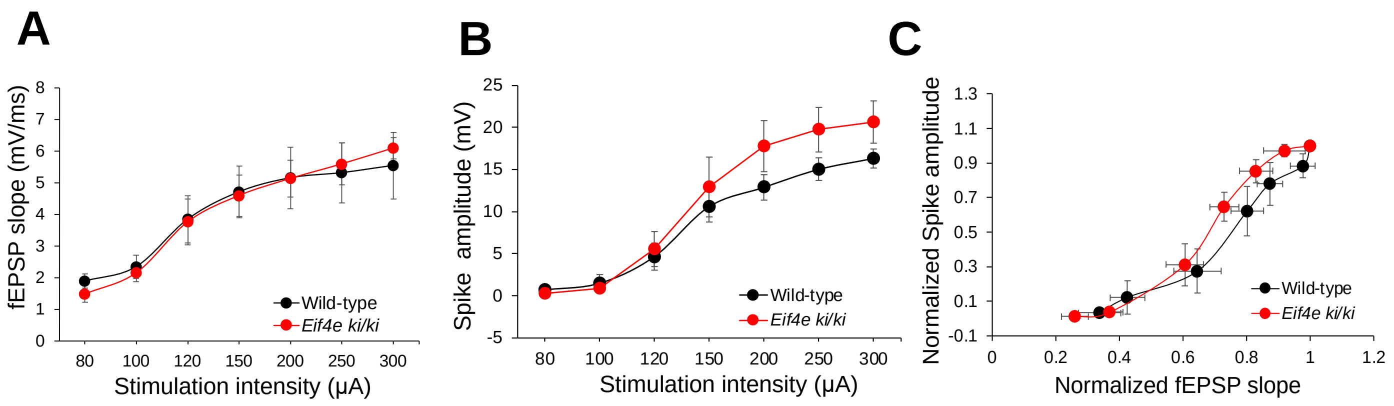

### Supp Fig S2

#### Supplemental Figure S2

A

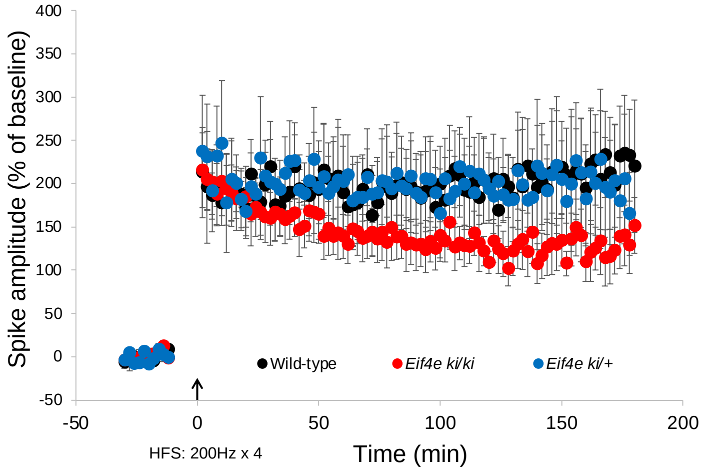


B

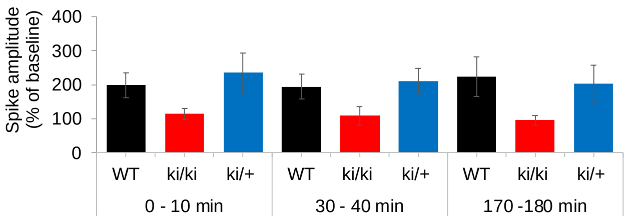

### Supp Fig S3

#### Supplemental Figure S3

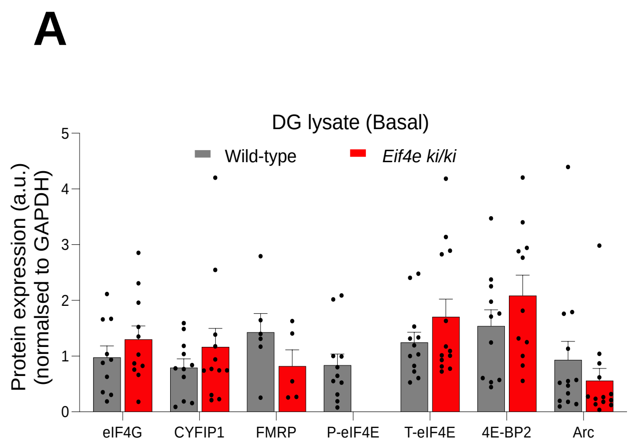


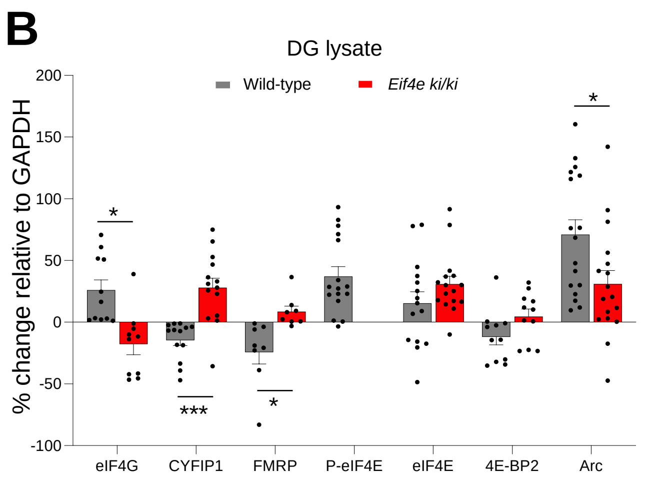


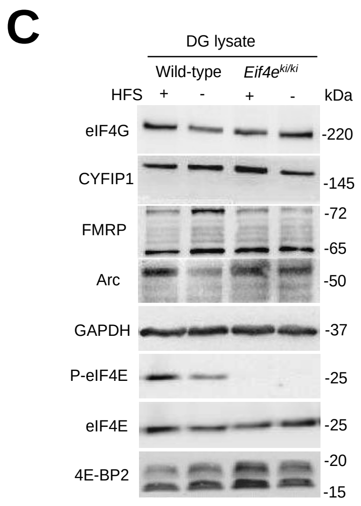

### Supp Fig S4

#### Supplemental Figure S4

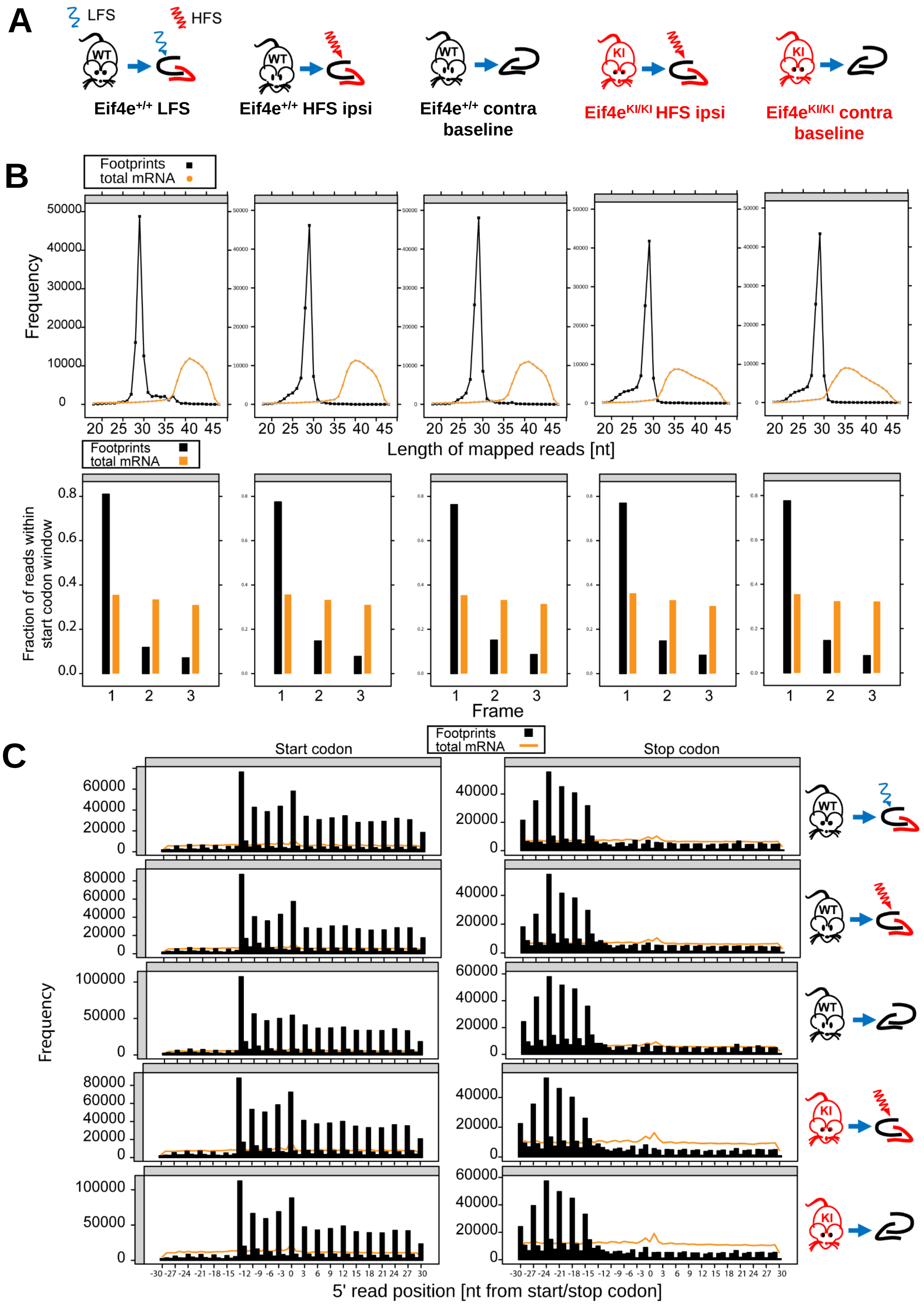

### Supp Fig S6

#### Supplemental Figure S6

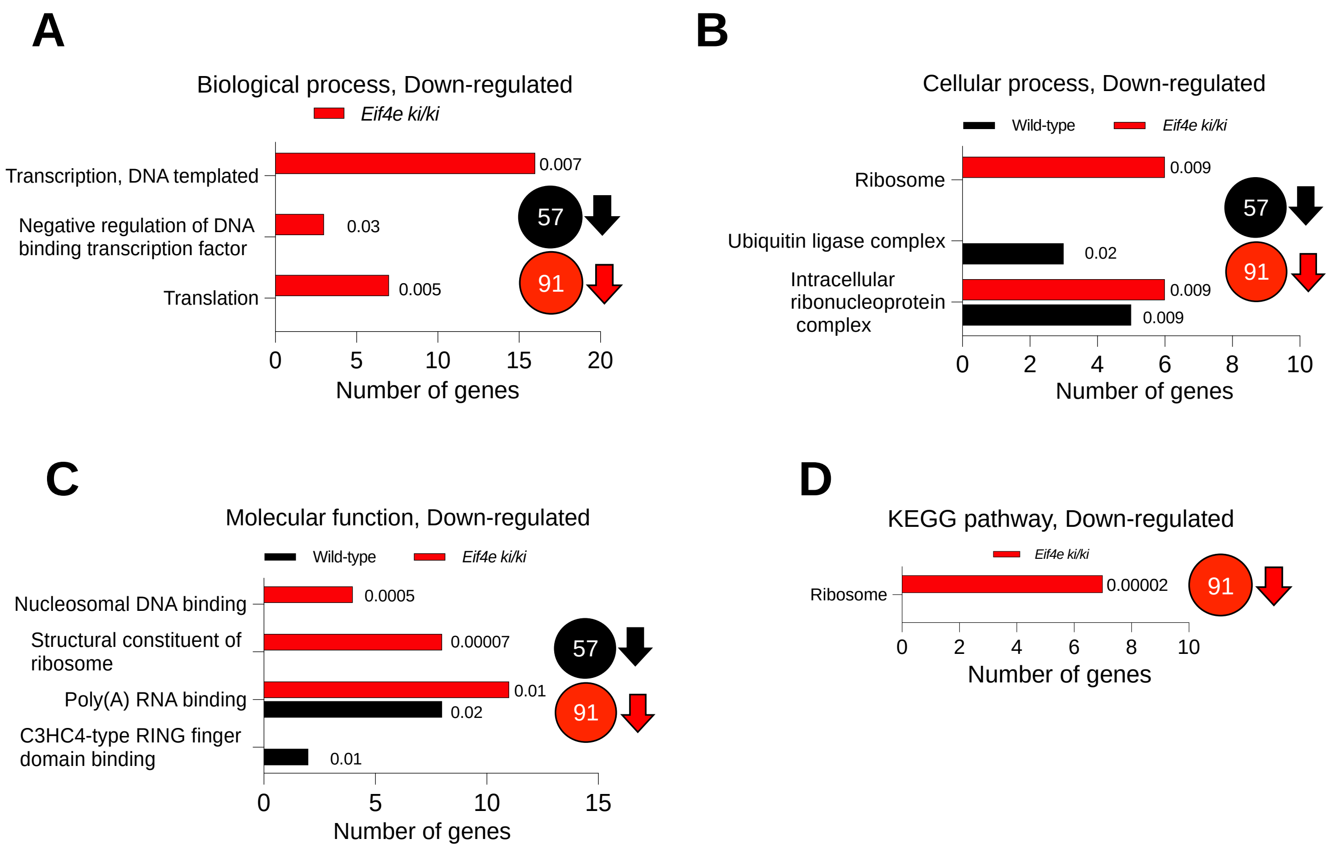

### Supp Fig S8

#### Supplemental Figure S8

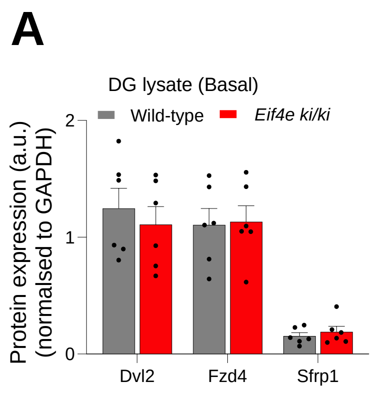


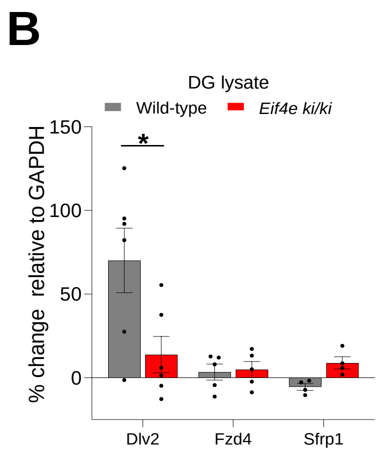


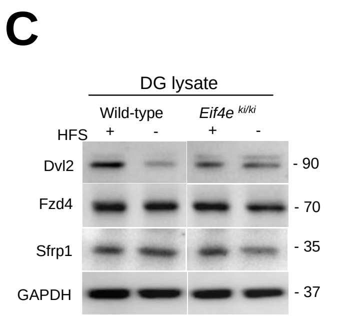
