## Supplementary material for "eIF4E phosphorylation recruits β-catenin to mRNA cap and selectively promotes Wnt pathway translation in dentate gyrus LTP maintenance in vivo": Supp Fig S5

### Supplemental Figure S5

A

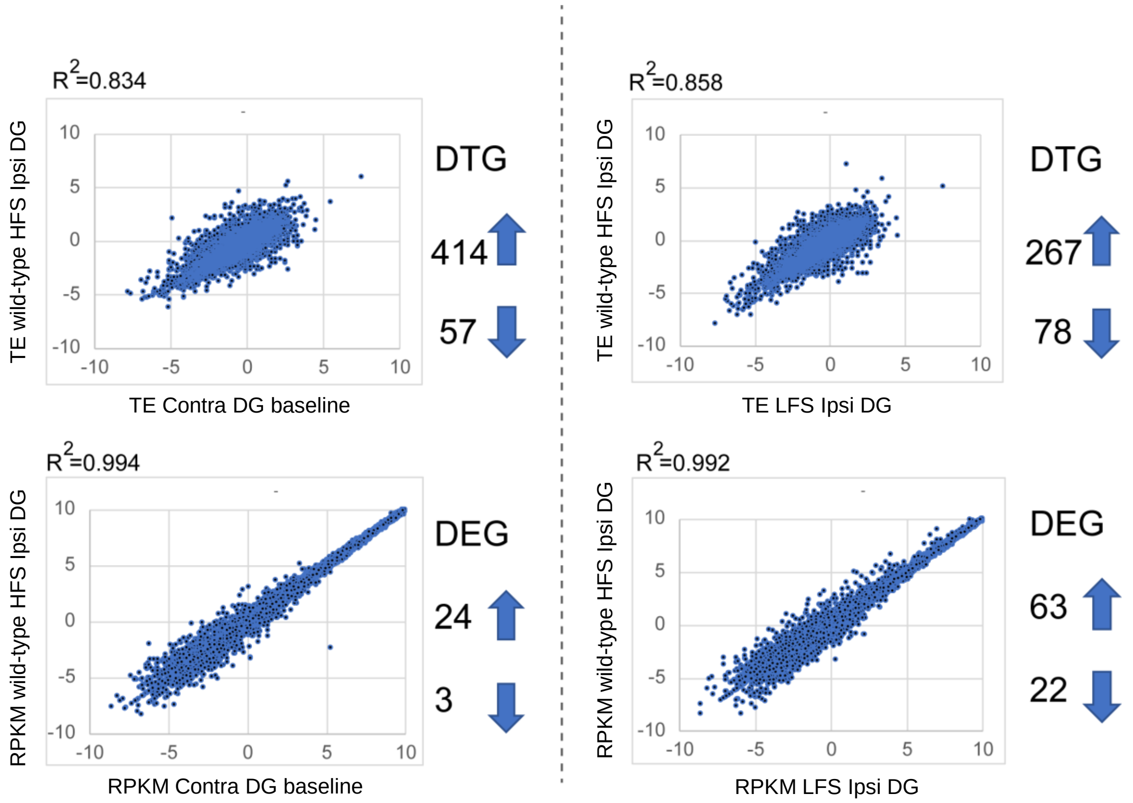


B

| | DTG IPA Canonical pathways | DEG IPA Canonical pathways |
|---|---|---|
| **Wild-type HFS Ipsi DG vs Contra DG** | 14-3-3-mediated Signaling | GP6 Signaling Pathway |
| | AMPK Signaling | Hepatic Fibrosis/Hepatic Stell. Cell Act. |
| | Actin Cytoskeleton Signaling | Growth Hormone Signaling |
| **Wild-type HFS Ipsi DG vs LFS Ipsi DG** | AMPK Signaling | Glucocorticoid Receptor Signaling |
| | Actin Nucleation by ARP-WASP Complex | Neuroinflammation Signaling Pathway |
| | Agranulocyte Adhesion and Diapedesis | Sirtuin Signaling Pathway |
