## Supplementary material for "eIF4E phosphorylation recruits β-catenin to mRNA cap and selectively promotes Wnt pathway translation in dentate gyrus LTP maintenance in vivo": Supp Fig S7

### Supplemental Figure S7

**A**

| | 5’ UTR analysis DTGs DOWN vs UP | |
|---|---|---|
| Feature | WT 40 min HFS | KI 40 min HFS |
| length | * | X |
| GC content | *** | * |
| Gibbs Free energy | *** | X |

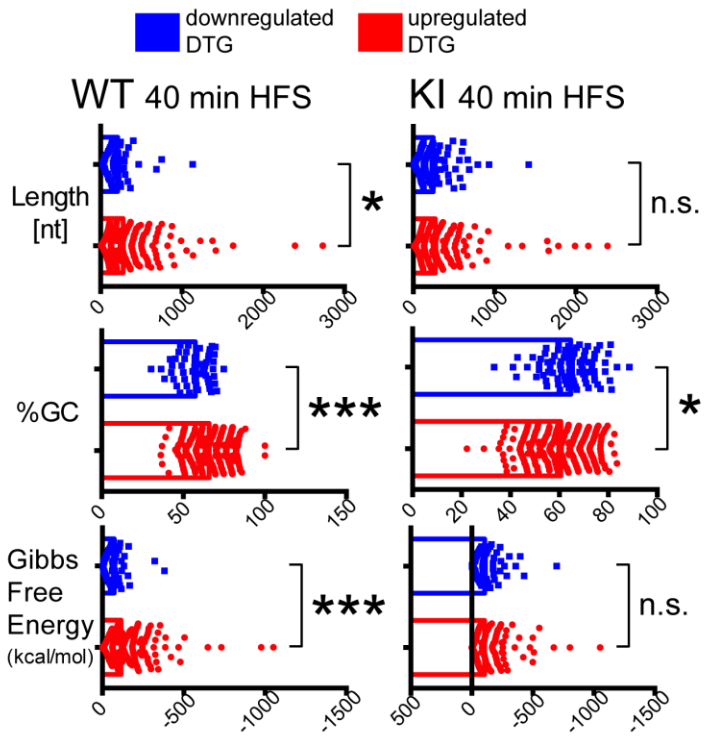


**B**

| | 5’ UTR motifs DTGs DOWN vs UP | |
|---|---|---|
| Feature | WT 40 min HFS | KI 40 min HFS |
| uORF | * | X |
| TOP | * | X |
| IRES | X | * |
| PG4 | * | X |

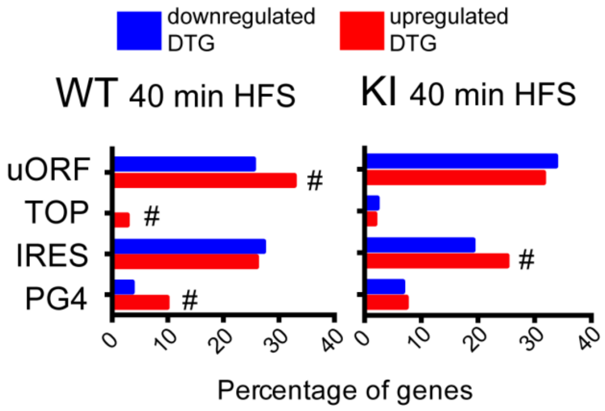
