## Supplementary material for "eIF4E phosphorylation recruits β-catenin to mRNA cap and selectively promotes Wnt pathway translation in dentate gyrus LTP maintenance in vivo": Supp Fig S9

### Supplemental Figure S9

**A**

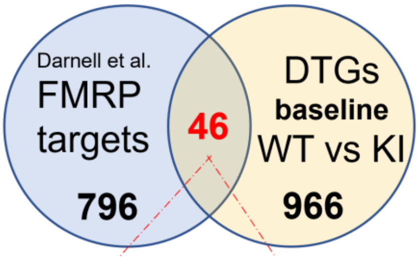


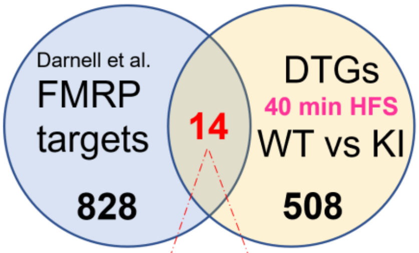


| | |
|---|---|
| Shank1 | Celsr3 |
| Kif5a | Zc3h7b |
| Nlgn2 | Lmtk3 |
| Lingo1 | Zfhx2 |
| Grik5 | Hdac5 |
| Zmiz2 | Tshz1 |
| Dlgap3 | Dtx1 |
| Ski | Lrrn2 |
| Gm996 | Shank2 |
| Lars2 | Kcnc3 |
| Cdk5r2 | Bcl9l |
| Prr12 | Pkd1 |
| Cbx6 | Mfhas1 |
| Atn1 | Tmem132a |
| Bcr | Nlgn3 |
| Chst2 | Crtc1 |
| Igsf9b | Hdac4 |
| Ttbk2 | C2cd2l |
| Psd | Dot1l |
| Syt7 | B3gat1 |
| Hivep3 | Synpo |
| Bzrap1 | Jph4 |
| Ttyh3 | Camk2n1 |

| |
|---|
| Pip5k1c |
| Git1 |
| Kcnb1 |
| Ckb |
| Zfp704 |
| Calm3 |
| Hcn2 |
| Ppp1r9b |
| Kcnc3 |
| Lrp3 |
| Fscn1 |
| Tubb3 |
| Agpat3 |
| Grm4 |

**B**

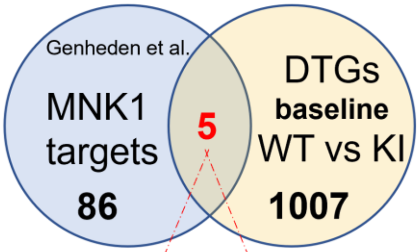


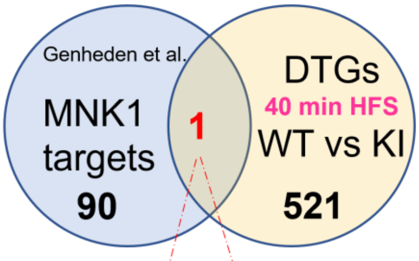


| |
|---|
| Sfrp1 |
| Gps1 |
| Atat1 |
| Kif5a |
| Fosb |

| |
|---|
| Flna |
