## Supplementary material for "eIF4E phosphorylation recruits β-catenin to mRNA cap and selectively promotes Wnt pathway translation in dentate gyrus LTP maintenance in vivo": Supp Table S1

Supplementary Table 1. List of Antibodies used.

| Antibody | Species | Source | Dilution |
| --- | --- | --- | --- |
| Arc | Mouse | #sc-17839; Santacruz biotechnology | 1:500 |
| CYFIP1 | Rabbit | #07-531; Upstate technology | 1:1000 |
| FMRP | Rabbit | #17722; Abcam | 1:1000 |
| 4E-BP2 | Rabbit | #2845; Cell signalling | 1:1000 |
| GAPDH | Mouse | #sc-32233; Santacruz biotechnology | 1:5000 |
| P-eIF4E | Rabbit | #9741; Cell signalling | 1:1000 |
| eIF4E | Rabbit | #9742; Cell signalling | 1:1000 |
| eIF4G | Rabbit | #2498; Cell signalling | 1:1000 |
| p-MNK1 Thr197/202 | Rabbit | #2111; Cell signalling | 1:1000 |
| MNK1 | Rabbit | #2195; Cell signalling | 1:1000 |
| 4E-BP1 Thr37/46 | Rabbit | #2855; Cell signalling | 1:1000 |
| RPL13a | Rabbit | #2765; Cell signalling | 1:500 |
| EPRS | Rabbit | #31531; Abcam | 1:3000 |
| Total-β-catenin | Rabbit | #19022; Millipore | 1:5000 |
| P-β-catenin S552 | Rabbit | #5651; Cell signalling | 1:1000 |
| P-β-catenin S33/T41 | Rabbit | #9561; Cell signalling | 1:1000 |
| Active-β-catenin S33/T41 | Rabbit | #05-665; Millipore | 1:1000 |
| Anti-Fzd4 | Rabbit | #ASJ-YDM4RO Nordic Biosite | 1:1000 |
| Sfrp1 | Rabbit | #16304905 Fisher scientific | 1:1000 |
| Dvl2 | Rabbit | #111532370 Fisher scientific | 1:1000 |
