## Supplementary material for "eIF4E phosphorylation recruits β-catenin to mRNA cap and selectively promotes Wnt pathway translation in dentate gyrus LTP maintenance in vivo": Supp Table S2

**Supplementary Table 2. Details of statistical analysis.**

| Figure | **Panel** | **mean± SEM** | **Group size** | **Statistics** | **Comparison** | **P-value** |
| --- | --- | --- | --- | --- | --- | --- |
| **1** | A |  |  |  |  |  |
|  | B | **Wild-type Vs. *Eif4e^Ki/Ki^***  30 min= 26.9 Vs 10.6  120 min= 29.1 Vs 5.6  180 min= 28.9 Vs 3.4 | Wild-type = 10  *Eif4e^Ki/Ki^* = 8 | Student  t-test. |  | **Wild-type Vs. *Eif4e^Ki/Ki^***  30 min= 0.00037  120 min= 0.00016  180 min= 2.08^-05^ |
|  |  | **Wild-type** = 26.9±10.6  *Eif4e^Ki/Ki^* = 16.3±3.0 | Wild-type =10  *Eif4e^KI/+^* = 7 | Student  t-test. |  | **Wild-type Vs. *Eif4e^KI/+^***  30 min= 0.022  120 min= 0.6393  180 min= 0.9089 |
| **2** | B | **Wild-type Vs. *Eif4e^Ki/Ki^***  P-MNK1/2 = 0.5Vs 0.7  T-MNK1/2 = 1.8 Vs 1.5  P-ERK1/2 = 2.9 Vs 2.3 T-ERK1/2 = 1.7 Vs 1.5 | P-MNK1/2 = 7  T-MNK1/2 = 11  P-ERK1/2 = 6  T-ERK1/2 = 6 | Multiple t-test. | Holm-Sidak  Method | **Wild-typeVs. *Eif4e^Ki/Ki^***  P-MNK1/2= 0.4159  T-MNK1/2= 0.7122  P-ERK1/2 = 0.3978  T-ERK1/2 = 0.5933 |
|  | C | **Wild-type Vs. *Eif4e^Ki/Ki^***  eIF4G = 0.61 Vs 0.59  CYFIP1= 0.50 Vs 0.53  FMRP= 0.82 Vs 0.74  T-BP2= 1.05 Vs 1.39 | eIF4G = 13  CYFIP1= 11 FMRP = 8  T-BP2 = 11 | Multiple t-test. | Holm-Sidak  Method | **Wild-type Vs. *Eif4e^Ki/Ki^***  eIF4G = 0.8907  CYFIP1= 0.8650  FMRP= 0.7343  T-BP2= 0.1518 |
|  | D | **Wild-type Vs. *Eif4e^Ki/Ki^***  P-/T-ERK1/2 = 86.8  P-/T-ERK1/2 = 64.93 | Wild-type = 6  *Eif4e^Ki/Ki^* = 6 | Student  t-test. | Holm-Sidak  Method | **Wild-type Vs. *Eif4e^Ki/Ki^***  P-/T-ERK1/2= 0.7507 |
|  | E | **Wild-typeVs. *Eif4e^Ki/Ki^***  P-/T-MNK1 = 26.2  P-/T-MNK1 = 24.1 | P-MNK1 = 12  T-MNK1 = 10 | Student  t-test. | Holm-Sidak  Method | **Wild-type Vs. *Eif4e^Ki/Ki^***  P-/T-MNK1 = 0.5026 |
|  | F | **Wild-typeVs. *Eif4e^Ki/Ki^***  eIF4G = 54.5 Vs -21.5  CYFIP1= -43.1 Vs 29.4  FMRP= -27.6 Vs 23.0  T-BP2= -14.7 Vs 31.4 | eIF4G = 13  CYFIP1= 12 FMRP= 7  T-BP2= 11 | Multiple t-test. | Holm-Sidak  Method | **Wild-type Vs. *Eif4e^Ki/Ki^***  eIF4G = 0.00017  CYFIP1= 0.00004  FMRP= 0.008  T-BP2= 0.0005 |
| **4** | A | **Wild-type Vs. *Eif4e^Ki/Ki^***  P-^S552^ β-cat = 0.46 Vs 0.45 Total- β-cat = 1.0 Vs 0.93  P-^~~S33/T41~~^β-cat = 0.26 Vs 0.43  P-^S33/T41^β-cat = 0.49 Vs 0.48 | P-^S552^β-cat = 12  Total-β-cat = 9  P-^~~S33/T41~~^β-cat =10  P-^S33/T41^β-cat = 9 | Multiple t-test. | Holm-Sidak  Method | **Wild-type Vs. *Eif4e^Ki/Ki^***  P-^S552^ β-cat = 0.9503  Total- β-cat = 0.7482  P-^~~S33/T4~~1^ β-cat = 0.2510  P-^S33/T41^ β-cat = 9.9314 |
|  | B | **Wild-type Vs. *Eif4e^Ki/Ki^***  P-^S552^ β-cat = 0.37 Vs 0.40  Total- β-cat = 0.56 Vs 0.47 | P-^S552^β-cat = 12  Total-β-cat = 12 | Multiple t-test. | Holm-Sidak  Method | **Wild-type Vs. *Eif4e^Ki/Ki^***  P-^S552^ β-cat = 0.8053  Total- β-cat = 0.6198 |
|  | D | **Wild-type Vs. *Eif4e^Ki/Ki^***  P-^S552^ β-cat = 26.0 Vs 20.9  Total- β-cat = 23.4 Vs 6.5  P-^~~S33/T41~~^β-cat = 47.0 Vs 27.8  P-^S33/T41^β-cat = 17.4 Vs 6.1 | P-^S552^ β-cat = 12  Total- β-cat = 9  P-^~~S33/T41~~^β-cat =10  P-^S33/T41^β-cat = 9 | Multiple t-test. | Holm-Sidak  Method | **Wild-type Vs. *Eif4e^Ki/Ki^***  P-^S552^ β-cat = 0.8230  Total- β-cat = 0.4362  P-^~~S33/T4~~1^ β-cat = 0.3847  P-^S33/T41^ β-cat = 0.5730 |
|  | E | **Wild-type Vs. *Eif4e^Ki/Ki^***  Total- β-cat = 78.3 Vs -13.3  P-^S552^ β-cat = 81.2 Vs 11.8  P-^S552^ /Total- β-cat=101.8Vs 114.4 | Total- β-cat = 12  P-^S552^ β-cat = 12  N=6 | Multiple t-test. | Holm-Sidak  Method | **Wild-type Vs. *Eif4e^Ki/Ki^***  Total- β-cat = 0.0013  P-^S552^ β-cat = 0.0074  P-^S552^ /Total-β-cat =0.7040 |
| **S1** | A | **Wild-type Vs. *Eif4e^Ki/Ki^*** 1. 1.889 Vs \| 1.484; 2. 2.344 Vs \| 2.152; 3. 3.842 Vs \| 3.759; 4. 4.707 Vs \| 4.589; 5. 5.332 Vs \| 5.127; 6. 5.634 Vs \| 5.588; 7. 5.863 Vs \| 6.088 | Wild-type= 4  *Eif4e^KI/Ki^* = 4 | Two-way  ANOVA |  | **Wild-type Vs. *Eif4e^Ki/Ki^***  0.9994 |
|  | B | **Wild-type Vs. *Eif4e^Ki/Ki^*** 1. 0.7027 Vs \| 0.2753; 2. 1.482 Vs \| 0.8794; 3. 4.642 Vs \| 5.533; 4. 10.59 Vs \| 12.95; 5. 12.89 Vs \| 17.78; 6. 15.03 Vs \| 19.74; 7. 16.33 Vs \| 20.61 | Wild-type= 4  *Eif4e^KI/Ki^* = 4 | Two-way  ANOVA |  | **Wild-type Vs. *Eif4e^Ki/Ki^***  0.5977 |
|  | C | **Wild-type Vs. *Eif4e^Ki/Ki^*** | Wild-type= 4  *Eif4e^KIKi^* = 4 | Three-way  ANOVA |  | **Wild-type Vs. *Eif4e^Ki/Ki^***  0.7631 |
| S2 | B | **Wild-type Vs. *Eif4e^Ki/Ki^***  30 min= 198.6 Vs 114.1  120 min= 194.2 Vs 108.6  180 min= 223.0 Vs 96.0 | Wild-type= 6  *Eif4e^Ki/Ki^* = 6 | Student  t-test. |  | **Wild-type Vs. *Eif4e^Ki/Ki^***  30 min= 0.062  120 min= 0.091  180 min= 0.063 |
| S3 | A | **Wild-type Vs. *Eif4e^Ki/Ki^*** eIF4G = 0.97 Vs 1.32  CYFIP1= 0.78 Vs 1.2  FMRP= 1.43 Vs 0.82  T-eIF4E= 1.25 Vs 1.7  T-BP2= 1.55 Vs 2.1  Arc = 0.94 Vs 0.56 | eIF4G = 10  CYFIP1= 12 FMRP = 6  T-eIF4E =13  T-BP2 = 12  Arc = 13 | Multiple t-test. | Holm-Sidak  Method | **Wild-type Vs. *Eif4e^Ki/Ki^***  eIF4G = 0.3234  CYFIP1= 0.3361  FMRP= 0.2139  T-eIF4E= 0.2319  T-BP2= 0.2537  Arc= 0.3545 |
|  | B | **Wild-typeVs. *Eif4e^Ki/Ki^***  eIF4G = 26.0 Vs -9.1  CYFIP1= -14.6 Vs 27.9  FMRP= -24.3 Vs 8.5  T-eIF4E= 25.0 Vs 20.6  P-BP2= 39.6 Vs 22.0  T-BP2= -11.9 Vs 4.6  Arc = 71.0 Vs 31.0 | eIF4G = 12  CYFIP1= 14 FMRP= 8  T-eIF4E=16  P-BP2= 6  T-BP2= 11  Arc= 17 | Multiple t-test. | Holm-Sidak  Method | **Wild-type Vs. *Eif4e^Ki/Ki^***  eIF4G = 0.02234  CYFIP1= 0.00007  FMRP= 0.0072  T-eIF4E= 0.7712  P-BP2= 0.0810  T-BP2= 0.0756  Arc= 0.018 |
| S7 | A | **Wild-type (Up vs Down)**  5´UTR length – 218.5 vs 160.0  5´UTR GC content – 65.96 vs 57.95  5´UTR Gibbs free energy –  -85.10 vs -55.95 | N= 404 vs 55 | Mann-Whitney test |  | **Wild-type**  5´UTR length = 0.0132  5´UTR GC content = 0.0001  5´UTR Gibbs free energy = 0.0003 |
|  | B | ***Eif4e^Ki/Ki^*** **(Up vs Down)**  5´UTR length – 200.5 vs 176.0  5´UTR GC content - 60.99 vs 64.86  5´UTR Gibbs free energy –  -69.15 vs -74.80 | N= 326 vs 89 |  |  | ***Eif4e^Ki/Ki^***  5´UTR length = 0.2170  5´UTR GC content = 0.0020  5´UTR Gibbs free energy = 0.9569 |
| S8 | A | **Wild-type Vs. *Eif4e^Ki/Ki^***  Dvl2 = 1.24 Vs 1.1  Fzd4 = 1.10 Vs 1.13  Sfrp1= 0.15 Vs 0.19 | Dvl2 = 6  Fzd4= 6  Sfrp1 = 6 | Multiple t-test. | Holm-Sidak  Method | **Wild-type Vs. *Eif4e^Ki/Ki^***  Dvl2 = 0.5651  Fzd4= 0.8947  Sfrp1= 0.5171 |
|  | B | **Wild-type Vs. *Eif4e^Ki/Ki^***  Dvl2 = 70.1 Vs 13.8  Fzd4 = 3.44 Vs 4.9  Sfrp1= -5.50 Vs 8.87 | Dvl2 = 6  Fzd4= 5  Sfrp1 = 5 | Multiple t-test. | Holm-Sidak  Method | **Wild-type Vs. *Eif4e^Ki/Ki^***  Dvl2 = 0.0295  Fzd4= 0.8320  Sfrp1= 0.0142 |
